# A protein kinase D–caveolin axis drives secretory carrier biogenesis at the trans-Golgi network

**DOI:** 10.64898/2026.01.13.699385

**Authors:** Hideaki Sugiura, Megumi Fujii, Yugo Terashima, Yuhei Takagi, Jessica Angulo-Capel, Eugenia Almacellas, Mitsuo Tagaya, Hiroki Inoue, Kohei Arasaki, Felix Campelo, Yuichi Wakana

**Author notes:** Corresponding author: Yuichi Wakana. H. Sugiura, M. Fujii, Y. Terashima, and Y. Takagi contributed equally to this paper.

## Abstract

Constitutive secretion from the trans-Golgi network (TGN) to the cell surface proceeds via carriers thought to form without a canonical cytoplasmic coat, yet how these carriers are generated remains poorly understood. Here, we identify a distinct population of TGN-to-cell surface carriers transporting influenza hemagglutinin (HA) and uncover a coat-like mechanism underlying their formation. HA carrier biogenesis requires non-vesicular lipid transfer at endoplasmic reticulum (ER)-Golgi membrane contact sites and protein kinase D (PKD) activity. We show that the integral membrane protein caveolin promotes membrane budding by assembling into cholesterol- and PKD-associated oligomers that act as a membrane-embedded, coat-like scaffold at lipid nanodomain-enriched TGN subdomains. These findings establish caveolin as a structural and regulatory component of TGN export and support a model in which a PKD–caveolin axis couples ER–Golgi lipid transfer to cargo sorting, membrane remodeling and fission during secretory carrier biogenesis.

## Introduction

Intracellular membrane trafficking depends on the continuous formation of transport carriers that mediate cargo exchange between organelles and the cell surface (Farquhar & Palade, 1981). The biogenesis of transport carriers is generally driven by cytoplasmic coat complexes: (1) clathrin-coated vesicles for endocytosis and Golgi-to-endosome transport (Pearse, 1975, 1976; Robinson, 2015); (2) COPI-coated vesicles for Golgi-to-endoplasmic reticulum (ER) and intra-Golgi transport (Malhotra *et al*, 1989; Serafini *et al*, 1991); and (3) COPII-coated vesicles for ER export (Barlowe *et al*, 1994). These cytoplasmic coats, together with their accessory proteins, couple cargo selection to membrane remodeling and fission (Kirchhausen, 2000). By contrast, constitutive transport from the trans-Golgi network (TGN) to the plasma membrane (PM) is thought to occur via pleiomorphic carriers that lack a canonical cytoplasmic coat. Despite their major role in secretion, the mechanisms underlying their biogenesis remain poorly understood (Stalder & Gershlick, 2020; Ford *et al*, 2021; Wakana & Campelo, 2021; Watson *et al*, 2025).

Protein kinase D (PKD) is a key regulator of TGN export (Liljedahl *et al*, 2001; Baron & Malhotra, 2002; Wakana & Campelo, 2021). PKD is recruited to the TGN by diacylglycerol (DAG), and its kinase activity is required for the fission of multiple classes of secretory carriers, including CARTS (carriers of the TGN to the cell surface), which transport selective cargoes such as pancreatic adenocarcinoma upregulated factor (PAUF), TGN46, and lysozyme C (Wakana *et al*, 2012). Subsequent studies demonstrated that CARTS biogenesis requires non-vesicular lipid transfer at ER–Golgi membrane contact sites (MCSs) (Wakana *et al*, 2015, 2021). At these MCSs, a complex of oxysterol-binding protein (OSBP) and vesicle-associated membrane protein-associated protein (VAP) transfers cholesterol from the ER to trans-Golgi/TGN membranes, coupled to the counter-transport of phosphatidylinositol 4-phosphate (PI4P) (Mesmin *et al*, 2013). PI4P is hydrolyzed by the ER-localized phosphatase Sac1, providing the driving force for cholesterol transfer (Antonny *et al*, 2018). In parallel, a ceramide transport protein (CERT) –VAP complex transports ceramide from the ER to the trans-Golgi/TGN membranes, where sphingomyelin (SM) synthases convert ceramide and phosphatidylcholine into SM and DAG (Hanada *et al*, 2003; Goto *et al*, 2020). DAG generated downstream of ceramide transfer recruits and activates PKD, most likely at sites adjacent to ER–Golgi MCSs, where PKD activity is essential for membrane fission during CARTS biogenesis (Wakana *et al*, 2015, 2021).

Lipid transfer at ER–Golgi MCSs is therefore central to establishing lipid gradients along the secretory pathway, particularly the enrichment of cholesterol and SM at trans-Golgi/TGN membranes. These lipids can assemble into liquid-ordered nanodomains that are depleted in other lipid species and these nanodomains may serve as platforms for selective protein recruitment (Simons & Ikonen, 1997; Sezgin *et al*, 2017). Increasing evidence indicates that such nanodomains may organize enzymatic activities, mediate cargo sorting, and promote the biogenesis of transport carriers, including CARTS (Duran *et al*, 2012; van Galen *et al*, 2014; Wakana *et al*, 2015; Campelo *et al*, 2017; Deng *et al*, 2018; Sundberg *et al*, 2019; Wakana *et al*, 2021; Anwar *et al*, 2022; Kovács *et al*, 2023; Castello-Serrano *et al*, 2024). Yet, the molecular machinery operating at these lipid nanodomains to drive budding and fission remains poorly defined.

To address this question, we established a synchronized transport system for influenza virus hemagglutinin (HA), whose exit from the TGN to the apical PM has been linked to its sorting into TGN lipid nanodomains (Scheiffele *et al*, 1997; Keller & Simons, 1998). This system enabled direct visualization of TGN-derived HA carriers, revealing that they exclude canonical cytoplasmic coats such as clathrin, COPI, and COPII, but are enriched in SM and thus potentially associated with lipid nanodomains. Similar to CARTS, HA carrier biogenesis depends on lipid transfer at ER–Golgi MCSs and on PKD activity. We further identified a functional and physical interaction between PKD and caveolin (CAV), a cholesterol-binding integral membrane protein best known as a structural component of caveolae at the PM. Whereas caveolae have well-established roles in mechanoprotection, signaling, lipid homeostasis, and endocytosis, the function of CAV proteins at the TGN has remained elusive (Parton & Simons, 2007; Cheng & Nichols, 2016; Parton, 2018). Our data suggest that cholesterol- and PKD-associated CAV oligomers, acting as a membrane-embedded, coat-like scaffold, promote the budding of secretory carriers at lipid nanodomain-enriched TGN regions. Together, these findings reveal a PKD–CAV axis that operates downstream of ER–Golgi lipid transfer to coordinate cargo sorting, membrane remodeling and fission during secretory carrier biogenesis at the TGN. Beyond defining a mechanistic module for TGN export, our work suggests a broader principle in which lipid-driven nanodomains and membrane-embedded scaffolds can substitute for classical cytoplasmic coats to drive coatless carrier formation, with implications for polarized secretion, viral assembly, and membrane trafficking-related diseases.

## Results

### Identification of TGN-derived HA carriers

Following influenza virus infection, HA is synthesized in the host ER and transported to the Golgi complex, where it has been proposed to partition into cholesterol- and SM-enriched nanodomains before exiting the TGN in transport carriers destined for the PM (apical PM in polarized cells) (Rindler *et al*, 1985; Skibbens *et al*, 1989; Scheiffele *et al*, 1997; Keller & Simons, 1998). To directly visualize TGN-derived HA-containing carriers (hereafter referred to as “HA carriers”) and monitor their biogenesis, we engineered a synchronized transport system using a reverse dimerization strategy, in which D/D solubilizer-induced disassembly of FKBP12-based FM4 domains synchronously releases proteins from the ER (Rivera *et al*, 2000). We fused a signal sequence, mKate2, and FM4 to the transmembrane domain-containing C-terminal region (residues 491–563) of H7N1 HA, generating the chimera mKate2-FM4-HA, which was stably expressed in HeLa cells (HA-HeLa cells). Upon addition of the D/D solubilizer, pre-existing mKate2-FM4-HA aggregates in the ER rapidly disassembled, triggering synchronized trafficking of HA to the Golgi complex (Fig. 1A,B). At 25 min after transport initiation, mKate2-FM4-HA was predominantly localized to the TGN and peripheral puncta (Fig. 1A, inset), whereas at 50 min it had reached the PM, where it remained at later time points. At 30 min, peripheral mKate2-FM4-HA puncta largely lacked clathrin, COPI, and COPII coats (mean colocalization frequencies ∼10% or lower; Fig. 1C,D) but showed robust colocalization with the SM reporter EQ-SM (Deng *et al*, 2016) (Fig. 1E), consistent with them being TGN-to-PM transport carriers devoid of conventional cytoplasmic coats but containing SM-rich lipid nanodomains. Quantitative analysis showed that ∼75% of HA-positive puncta were positive for EQ-SM, whereas only ∼25% of EQ-SM–positive puncta were HA-positive (Fig. 1F), indicating that HA carriers represent a subset of SM-rich transport carriers.

**Figure 1.**
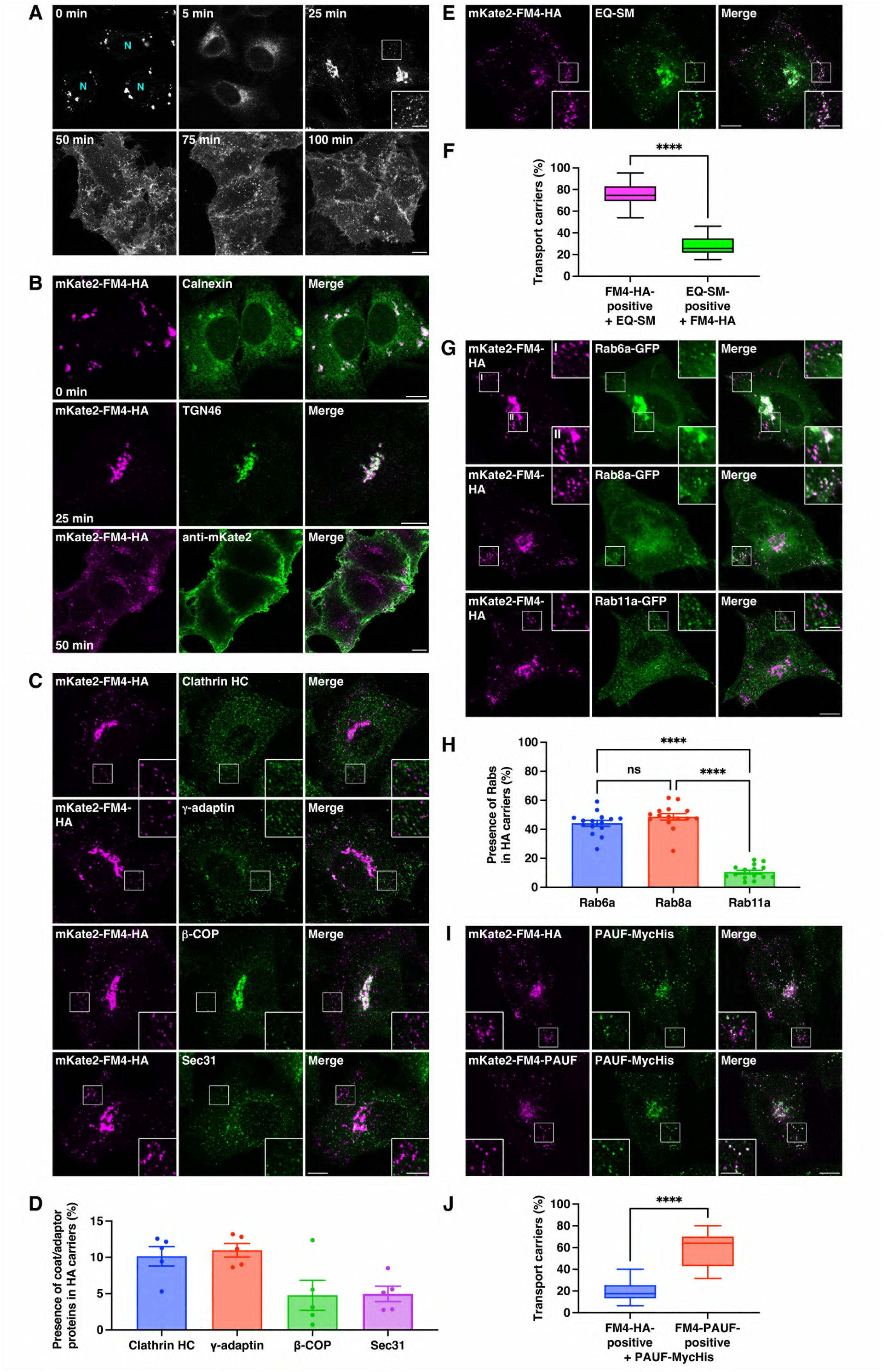
Identification of TGN-derived HA carriers. **(A)** mKate2-FM4-HA transport from the ER to the PM via the Golgi complex. HA-HeLa cells were incubated at 37°C with D/D solubilizer and cycloheximide, fixed at the indicated times, and visualized by mKate2 fluorescence. The inset shows a magnified image of the boxed area with brightness and contrast enhancement. N, nucleus. **(B)** Colocalization of mKate2-FM4-HA with the ER maker calnexin (0 min) and the TGN maker TGN46 (25 min) in permeabilized HA-HeLa cells and visualization of extracellularly located mKate2 (50 min) by labelling of non-permeabilized HA-HeLa cells with an anti-mKate2 antibody. **(C, D)** Distribution of HA carriers, clathrin-coated vesicles (labeled with anti-clathrin HC and anti-γ-adaptin antibodies), COPI-coated vesicles (labeled with an anti-β-COP antibody), and COPII-coated vesicles (labeled with an anti-Sec31 antibody) in HA-HeLa cells (30 min). **(D)** Quantification of the mKate2-FM4-HA– positive HA carriers containing clathrin HC, γ-adaptin, β-COP, or Sec31. Data are means ± SEM, individual points represent values from individual cells ([clathrin HC] mKate2-FM4-HA positive: n = 824 puncta and clathrin HC positive: n = 1,869 puncta in 5 cells; [γ-adaptin] mKate2-FM4-HA positive: n = 888 puncta and γ-adaptin positive: n = 1,896 puncta in 5 cells; [β-COP] mKate2-FM4-HA positive: n = 842 puncta and β-COP positive: n = 1,135 puncta in 5 cells; [Sec31] mKate2-FM4-HA positive: n = 928 puncta and Sec31 positive: n = 1,212 puncta in 5 cells). **(E, F)** Colocalization of HA carriers and EQ-SM. HA-HeLa cells expressing EQ-SM were incubated at 37°C for 30 min with D/D solubilizer and cycloheximide, fixed, and visualized by mKate2 fluorescence and oxGFP fluorescence from EQ-SM. **(F)** Quantification of the mKate2-FM4-HA–positive HA carriers containing EQ-SM (magenta) and EQ-SM–positive carriers containing mKate2-FM4-HA (green). Boxes delimit the first and third quartiles, and the central line is the median. The whiskers represent the minimum and maximum values (mKate2-FM4-HA positive: n = 1,631 puncta; EQ-SM positive: n = 4,055 puncta in 20 cells; ****, P < 0.0001; paired two-tailed Student’s t test). **(G, H)** Colocalization of HA carriers with Rab6a-GFP and Rab8a-GFP, but not with Rab11a-GFP. HA-HeLa cells expressing each Rab-GFP construct were incubated at 37°C for 30 min with D/D solubilizer and cycloheximide, fixed, and visualized by mKate2 and GFP fluorescence. Arrowheads indicate tubular structures containing both Rab6a-GFP and mKate2-FM4-HA. **(H)** Quantification of the mKate2-FM4-HA–positive HA carriers containing Rab6a-GFP, Rab8a-GFP, or Rab11a-GFP. Data are means ± SEM, individual points represent values from individual cells ([Rab6a] mKate2-FM4-HA positive: n = 1,574 puncta and Rab6a-GFP positive: n = 2,756 puncta in 15 cells; [Rab8a] mKate2-FM4-HA positive: n = 2,144 puncta and Rab8a-GFP positive: n = 6,340 puncta in 15 cells; [Rab11a] mKate2-FM4-HA positive: n = 1,974 puncta and Rab11a-GFP positive: n = 5,740 puncta in 15 cells; ****, P < 0.0001; one-way ANOVA multiple comparison test). **(I, J)** Distribution of HA carriers and CARTS. HeLa cells stably expressing PAUF-MycHis were incubated at 37°C for 30 min with D/D solubilizer and cycloheximide, fixed, and visualized by mKate2 fluorescence and anti-Myc immunostaining. **(J)** Quantification of the mKate2-FM4-HA–positive HA carriers containing PAUF-MycHis (blue) and mKate2-FM4-PAUF–positive CARTS containing PAUF-MycHis (red). Boxes delimit the first and third quartiles, and the central line is the median. The whiskers represent the minimum and maximum values (blue, mKate2-FM4-HA positive: n = 829 puncta and PAUF-MycHis positive: n = 870 puncta in 15 cells; red, mKate2-FM4-PAUF positive: n = 461 puncta and PAUF-MycHis positive: n = 603 puncta in 15 cells; ****, P < 0.0001; paired two-tailed Student’s t test). The insets in **(C, E, G, I)** show magnified images of the boxed areas. Scale bars, 10 µm (large panels) and 5 µm (insets). Source data are available online for this figure.

Because these features are shared with CARTS (Wakana *et al*, 2012, 2021), we next examined whether HA carriers also contained established trafficking machinery characteristic of CARTS. We analyzed Rab6a and Rab8a, two small GTPases associated with CARTS, together with Rab11a, which is predominantly associated with recycling endosomes and absent from CARTS (Wakana *et al*, 2012). At 30 min after transport initiation, HA carriers overlapped with Rab6a- and Rab8a-positive puncta, but not with Rab11a (Fig. 1G), consistent with previous observations of CARTS. In Rab6a-GFP–expressing cells, we additionally observed tubular structures containing both Rab6a and HA (Fig. 1G, arrowheads). Quantification showed that ∼40% of HA carriers were positive for Rab6a-GFP and for Rab8a-GFP (Fig. 1H), indicating that HA carriers share key trafficking machinery with CARTS. We next asked whether this similarity also extended to cargo composition. When the CARTS-specific cargo MycHis-tagged PAUF was co-expressed with either mKate2-FM4-HA or mKate2-FM4-PAUF as a positive control, only ∼18% of mKate2-FM4-HA–positive carriers contained PAUF-MycHis, whereas ∼65% overlap was observed in the PAUF control (Fig. 1I,J). Taken together, these results indicate that, although HA carriers share several components of the CARTS trafficking machinery, they differ in cargo composition and thus represent a distinct population of CARTS-like TGN-derived transport carriers.

### Lipid transfer at ER–Golgi MCSs is required for HA carrier biogenesis

Cholesterol in the TGN membranes has been reported to be important for the export of HA from the TGN (Keller & Simons, 1998), yet the source of this cholesterol has remained unclear. To test whether lipid transfer at ER–Golgi MCSs is required for HA transport, we depleted key components of ER–Golgi MCSs by siRNA-mediated knockdown. In HA-HeLa cells depleted of two isoforms of VAP (VAP-A and VAP-B), OSBP, or Sac1, synchronized transport of mKate2-FM4-HA from the ER to the Golgi complex was largely unaffected by 25 min post-release (Fig. 2A,B). By contrast, by 50 min, surface levels of HA were markedly reduced under each knockdown condition (Fig. 2A). In these cells, HA accumulated at the TGN (Fig. 2C), indicating a specific defect in TGN-to-PM trafficking. Similar phenotypes were obtained upon depletion of sterol regulatory element binding protein (SREBP) cleavage-activating protein (SCAP) –an ER cholesterol sensor that interacts with the OSBP–VAP complex through Sac1 to promote CARTS biogenesis (Wakana *et al*, 2021)– or upon depletion of CERT, albeit to a lesser extent (Fig. 2A–C). These results suggest that lipid transfer pathways at ER– Golgi MCSs contribute to HA export from the TGN.

**Figure 2.**
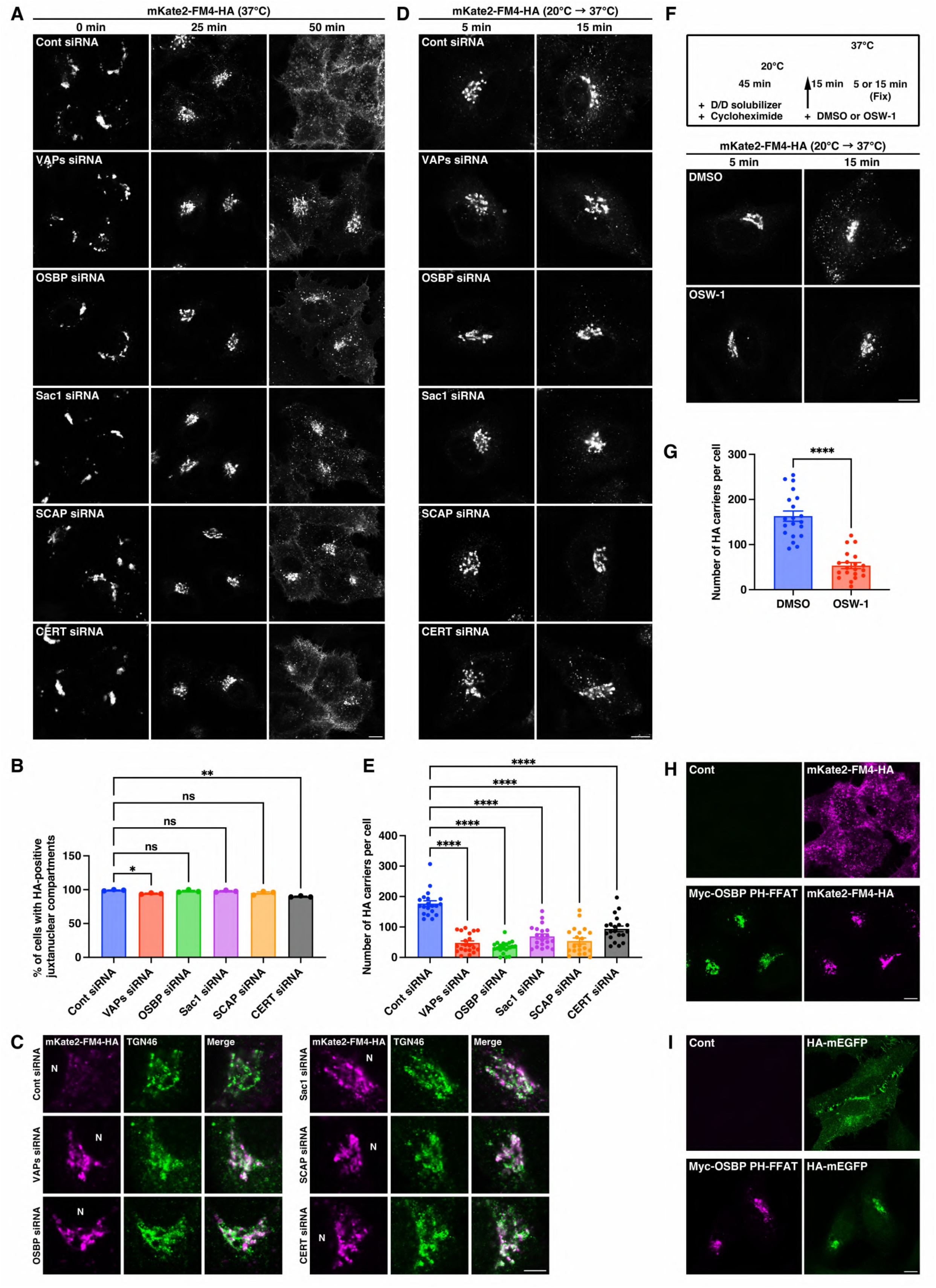
Lipid transfer at ER–Golgi MCSs is required for HA carrier biogenesis. **(A, B)** mKate2-FM4-HA transport from the ER to the PM via the Golgi complex under control (Cont), VAP-A/B (VAPs), OSBP, Sac1, SCAP, and CERT knockdown conditions in HA-HeLa cells. The cells were incubated at 37°C with D/D solubilizer and cycloheximide, fixed at the indicated times, and visualized by mKate2 fluorescence. Scale bar, 10 µm. **(B)** Quantification of mKate2-FM4-HA transport from the ER to the Golgi complex. The percentage of cells showing juxtanuclear accumulation of mKate2-FM4-HA at 25 min after transport initiation is shown. Data are means ± SEM (n = 3 independent experiments [102–134 cells per condition]; *, P = 0.0132; **, P = 0.0011; one-way ANOVA multiple comparison test). **(C)** Accumulation of mKate2-FM4-HA at the TGN under ER–Golgi MCS component knockdown conditions in HA-HeLa cells (50 min). The cells were fixed and visualized by mKate2 fluorescence and anti-TGN46 immunostaining. N, nucleus. Scale bar, 5 µm. **(D, E)** HA carrier biogenesis under control knockdown and ER–Golgi MCS component knockdown conditions in HA-HeLa cells. The cells were subjected to HA carrier formation assay as described in Methods, fixed, and visualized by mKate2 fluorescence. Scale bar, 10 µm. See also Movie EV1. **(E)** Quantification of the HA carrier biogenesis. The number of HA carriers at 15 min after the temperature shift to 37°C is shown. Data are means ± SEM (n = 20 cells per condition; ****, P < 0.0001; one-way ANOVA multiple comparison test). **(F, G)** HA carrier biogenesis in HA-HeLa cells treated with DMSO or 20 nM OSW-1. The cells were subjected to HA carrier formation assay as described in Methods, fixed, and visualized by mKate2 fluorescence. Scale bar, 10 µm. **(G)** Quantification of the HA carrier biogenesis. The number of HA carriers at 15 min after the temperature shift to 37°C is shown. Data are means ± SEM (n = 20 cells per condition; ****, P < 0.0001; unpaired two-tailed Student’s t test). **(H)** Accumulation of mKate2-FM4-HA at the Golgi complex due to Myc-OSBP PH-FFAT expression. Control HA-HeLa cells and those expressing Myc-OSBP PH-FFAT were incubated at 37°C for 50 min with D/D solubilizer and cycloheximide, fixed, and visualized by anti-Myc immunostaining and mKate2 fluorescence. Scale bar, 10 µm. **(I)** Accumulation of HA-mEGFP at the Golgi complex due to Myc-OSBP PH-FFAT expression. HeLa cells expressing HA-mEGFP alone (control) or co-expressing HA-mEGFP and Myc-OSBP PH-FFAT were fixed and visualized by anti-Myc immunostaining and mEGFP fluorescence. Scale bar, 10 µm. Source data are available online for this figure.

To directly assess HA carrier formation at the TGN, we employed a temperature block-and-release assay. Following accumulation of mKate2-FM4-HA in the TGN at 20°C for 45 min, cells were shifted back to 37°C to release the TGN export block, allowing for a TGN-synchronized and more efficient induction of HA carrier biogenesis. Live-cell imaging showed that the perinuclear HA signal progressively dispersed into numerous HA carriers destined to the cell surface (Movie EV1). Quantitative analysis showed that, by 15 min after release from the TGN, cells depleted of VAPs, OSBP, Sac1, SCAP, or, to a lesser extent, CERT exhibited a significant reduction in the number of HA carriers (Fig. 2D,E), consistent with the delayed TGN export observed under these conditions (Fig. 2A,C).

To further corroborate the role of cholesterol transfer at ER–Golgi MCSs in HA carrier biogenesis, we treated HA-HeLa cells with OSW-1, an OSBP inhibitor that blocks cholesterol transport from the ER to the trans-Golgi/TGN membranes (Mesmin *et al*, 2017). After a 20°C block for 45 min, cells were pretreated with dimethyl sulfoxide (DMSO) or OSW-1 at 20°C for additional 15 min before being shifted to 37°C. By 15 min after the temperature shift, OSW-1 treatment reduced the number of HA carriers to ∼33% of control levels (Fig. 2F,G), closely mirroring its effect on the biogenesis of mKate2-FM4-PAUF–containing CARTS (reduced to ∼31% of control levels; Fig. EV1A,B). Finally, expression in HA-HeLa cells of Myc-OSBP PH-FFAT, which stabilizes ER–Golgi MCSs while blocking OSBP-dependent cholesterol transport and CARTS biogenesis (Mesmin *et al*, 2013; Wakana *et al*, 2015), strongly inhibited TGN export of mKate2-FM4-HA (Fig. 2H), suggesting that dynamic assembly and disassembly of ER–Golgi MCSs is required for HA carrier biogenesis. Consistently, transport of full-length HA-mEGFP was similarly impaired in a non-synchronized transport assay (Fig. 2I). Taken together, these data indicate that lipid transfer at ER–Golgi MCSs is essential not only for CARTS formation but also for the biogenesis of HA carriers.

### PKD kinase activity is required for selective cargo sorting and membrane fission during HA carrier biogenesis

Ceramide transferred at ER–Golgi MCSs is used to produce DAG at the TGN, which in turn recruits PKD to promote carrier membrane fission (Peretti *et al*, 2008; Wakana *et al*, 2015; Wakana & Campelo, 2021). To assess the role of PKD in HA carrier formation, we knocked down the two PKD isoforms expressed in HeLa cells, PKD2 and PKD3 (Bossard *et al*, 2007), using two independent siRNAs per isoform. Immunoblotting confirmed efficient depletion of PKD2 and PKD3 relative to control siRNA (Appendix Fig. S1A). Notably, one PKD2-targeting siRNA (1303) also reduces PKD3 levels, likely owing to sequence homology. In HA-HeLa cells, PKD2 knockdown caused pronounced retention of mKate2-FM4-HA in the juxtanuclear Golgi region at 50 min after transport initiation, whereas PKD3 depletion resulted in only a modest delay (Fig. 3A). Consistent with this, HA carrier biogenesis was strongly inhibited upon PKD2 knockdown but only partially reduced following PKD3 depletion (Fig. 3B). Quantitative analysis showed that PKD2 depletion (siRNAs 882 and 1303) reduced the number of HA carriers to ∼11% and ∼13% of control levels, respectively, whereas PKD3 depletion (siRNAs 1363 and 3’) reduced carrier number to ∼57% and ∼66%, respectively (Fig. 3C). These data indicate a predominant requirement for PKD2, compared with PKD3, in HA carrier biogenesis in HeLa cells. Comparison with CARTS formation revealed that PKD2 knockdown similarly reduced the number of mKate2-FM4-PAUF–containing CARTS (Fig. 3D,E). However, under PKD2 depletion, PAUF but not HA was incorporated into elongated membrane tubules extending from the TGN, which likely represent fission-defective carrier precursors (Fig. 3D,F,G). These observations raise the possibility of PKD2 is additionally required for efficient sorting and loading of HA into nascent secretory carriers at the TGN.

**Figure 3.**
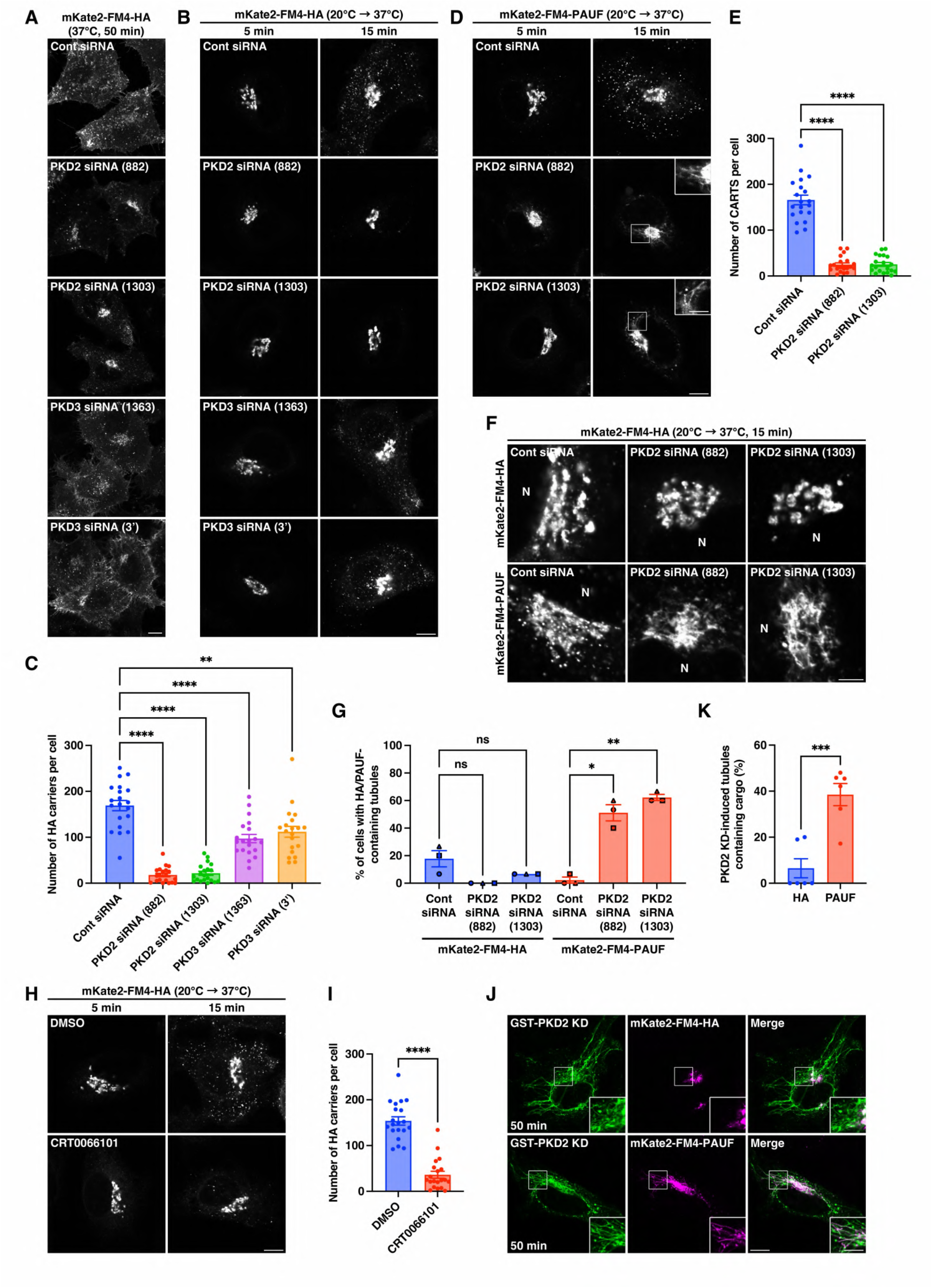
PKD kinase activity is required for selective cargo sorting and membrane fission during HA carrier biogenesis. **(A)** mKate2-FM4-HA transport to the PM under control (Cont), PKD2, and PKD3 knockdown conditions in HA-HeLa cells. The cells were incubated at 37°C for 50 min with D/D solubilizer and cycloheximide, fixed, and visualized by mKate2 fluorescence. Scale bar, 10 µm. **(B, C)** HA carrier biogenesis under control, PKD2, and PKD3 knockdown conditions in HA-HeLa cells. The cells were subjected to HA carrier formation assay as described in Methods, fixed, and visualized by mKate2 fluorescence. Scale bar, 10 µm. **(C)** Quantification of the HA carrier biogenesis. The number of HA carriers at 15 min after the temperature shift to 37°C is shown. Data are means ± SEM (n = 20 cells per condition; **, P < 0.005; ****, P < 0.0001; one-way ANOVA multiple comparison test). **(D, E)** CARTS biogenesis under control and PKD2 knockdown conditions in HeLa cells stably expressing mKate2-FM4-PAUF (PAUF-HeLa cells). The cells were subjected to CARTS formation assay as described in Methods, fixed, and visualized by mKate2 fluorescence. The insets show magnified images of the boxed areas with brightness and contrast enhancement. Scale bars, 10 µm (large panels) and 5 µm (insets). **(E)** Quantification of the CARTS biogenesis. The number of CARTS at 15 min after the temperature shift to 37°C is shown. Data are means ± SEM (n = 20 cells per condition; ****, P < 0.0001; one-way ANOVA multiple comparison test). **(F, G)** Distribution of mKate2-FM4-HA and mKate2-FM4-PAUF in the juxtanuclear Golgi region under control and PKD2 knockdown conditions in HA-HeLa and PAUF-HeLa cells. The cells were fixed and visualized by mKate2 fluorescence. N, nucleus. Scale bar, 5 µm. **(G)** Quantification of HA/PAUF incorporation into TGN tubules. The percentage of cells with HA/PAUF-containing tubules is shown. Data are means ± SEM (n = 3 independent experiments [15 cells per condition]; *, P = 0.0399; **, P = 0.0063; repeated-measures one-way ANOVA multiple comparison test for each cargo protein). **(H, I)** HA carrier biogenesis in HA-HeLa cells treated with DMSO or 5 μM CRT0066101. The cells were subjected to HA carrier formation assay as described in Methods, fixed, and visualized by mKate2 fluorescence. Scale bar, 10 µm. **(I)** Quantification of the HA carrier biogenesis. The number of HA carriers at 15 min after the temperature shift to 37°C is shown. Data are means ± SEM (n = 20 cells per condition; ****, P < 0.0001; unpaired two-tailed Student’s t test). **(J, K)** Distribution of mKate2-FM4-HA and mKate2-FM4-PAUF in HA-HeLa and PAUF-HeLa cells expressing GST-PKD2 KD. The cells were incubated at 37°C for 50 min with D/D solubilizer and cycloheximide, fixed, and visualized by anti-GST immunostaining and mKate2 fluorescence. The insets show magnified images of the boxed areas. Scale bars, 10 µm (large panels) and 5 µm (insets). **(K)** Quantification of HA/PAUF incorporation into GST-PKD2 KD-induced membrane tubules. The percentage of HA/PAUF-containing tubules is shown. Data are means ± SEM, individual points represent values from individual cells ([HA] mKate2-FM4-HA positive: n = 7 tubules and GST-PKD2 KD positive: n = 120 tubules in 6 cells; [PAUF] mKate2-FM4-PAUF positive: n = 57 tubules and GST-PKD2 KD positive: n = 153 tubules in 6 cells; ***, P = 0.0005; unpaired two-tailed Student’s t test). Source data are available online for this figure.

We next examined whether PKD2 depletion affects the steady-state localization of full-length HA using a non-synchronized transport assay. HA-mEGFP expression was induced for 24 h using a Tet-On system, 48 h after siRNA transfection. PKD2 knockdown significantly decreased the PM-to-Golgi fluorescence signal ratio of HA-mEGFP (Fig. EV2A,B), further supporting a role for PKD2 in full length HA trafficking.

Finally, pharmacological inhibition of PKD kinase activity with CRT0066101 significantly reduced the number of HA carriers (Fig. 3H,I), as well as of CARTS (Fig. EV1A,B). Similar to PKD2 depletion, CRT0066101 treatment induced the formation of TGN-associated tubules containing PAUF (Fig. EV1, inset) but not HA (Fig. 3H). Furthermore, expression of a dominant-negative, kinase-dead (KD) PKD2 mutant phenocopied these effects: both HA and PAUF accumulated in the juxtanuclear Golgi region, but only PAUF was clearly detected within the long TGN tubules induced by PKD2 KD expression (Fig. 3J,K). These results suggest that PKD2 kinase activity is necessary for efficient HA sorting and loading into nascent carriers, whereas PAUF sorting depends on the cargo receptor TGN46 but does not require PKD2 kinase activity (Lujan *et al*, 2024). Collectively, these results indicate that PKD2 kinase activity is essential for the biogenesis of both CARTS and HA carriers, and further suggest that, unlike CARTS, HA carriers require PKD2 not only for membrane fission but also for efficient cargo sorting and incorporation into nascent carriers at the TGN.

### PKD kinase activity is required for apical transport in polarized MDCK cells

Previous studies have implicated PKD in basolateral trafficking in polarized MDCK cells (Yeaman *et al*, 2004). Our observation that PKD kinase activity is required for efficient transport of the apical cargo HA in non-polarized HeLa cells prompted us to test whether PKD activity is similarly required for HA transport in polarized MDCK cells. To this end, we generated an MDCK cell line stably expressing mKate2-FM4-HA (HA-MDCK cells), which was cultured on Transwell filters to establish polarity. Upon addition of the D/D solubilizer, mKate2-FM4-HA was synchronously transported from the ER (0 min) to the Golgi complex (30 min) and subsequently to the apical membrane (60 min), where it formed progressively larger clusters by 90 min (Fig. EV3A,B). Such clustering was not apparent in HeLa cells (Fig. 1A), likely reflecting differences in apical surface morphology and organization between polarized MDCK cells and non-polarized HeLa cells.

To examine the effect of PKD inhibition in apical HA transport, polarized HA-MDCK cells were first incubated at 20°C for 45 min and treated with DMSO or CRT0066101 at 20°C for additional 15 min before being shifted to 37°C for 5 or 30 min. In DMSO-treated control cells, mKate2-FM4-HA largely remained in the Golgi membranes at 5 min and reached the apical PM by 30 min (Fig. 4A, left panels). By contrast, CRT0066101-treated cells retained the majority of HA in the Golgi membranes even at 30 min (Fig. 4A, right panels). Quantitative analysis revealed a significant reduction in the apical PM-to-Golgi fluorescence signal ratio of mKate2-FM4-HA upon PKD inhibition (Fig. 4B).

**Figure 4.**
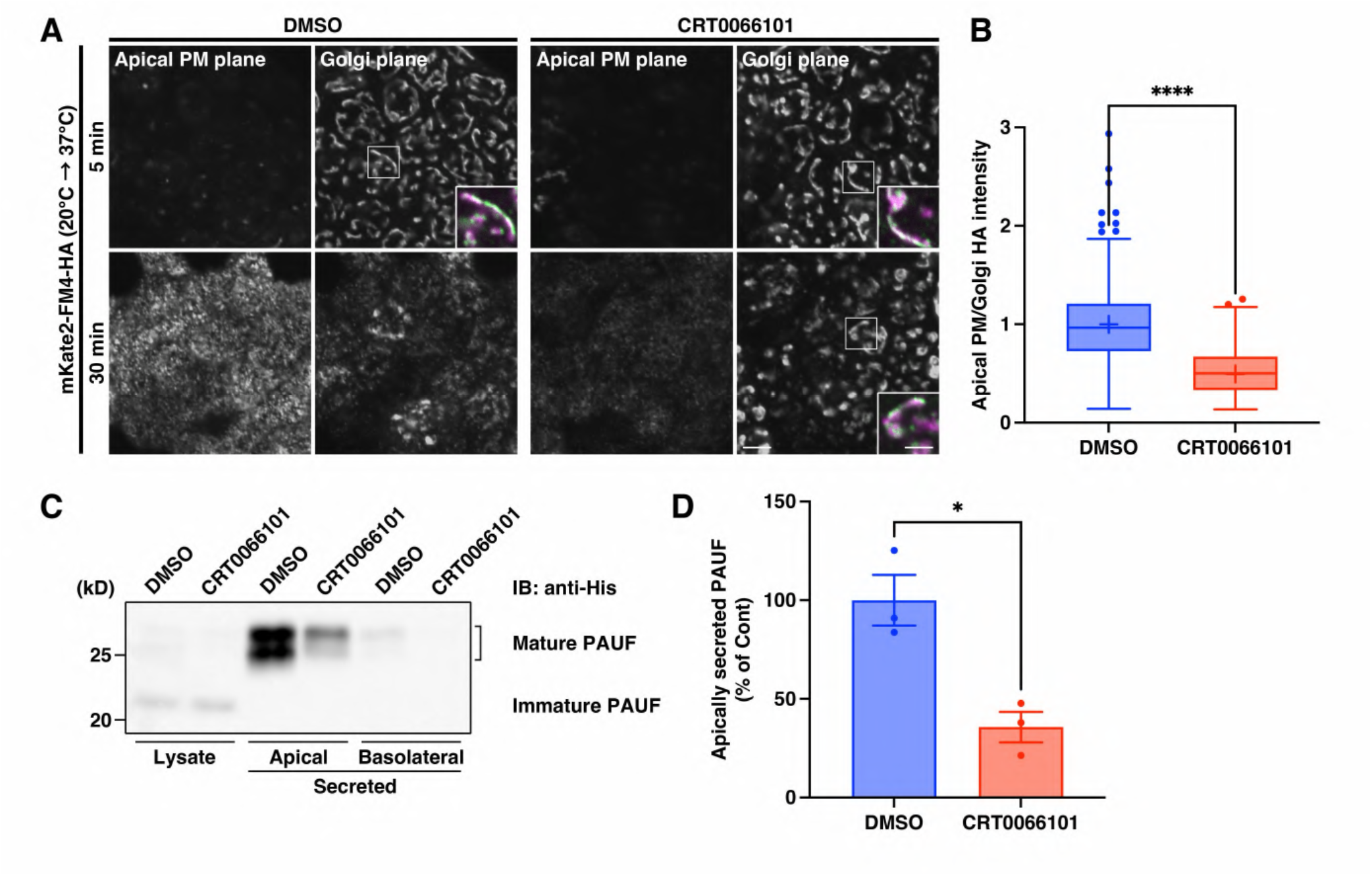
PKD kinase activity is required for apical transport in polarized MDCK cells. **(A, B)** mKate2-FM4-HA transport from the Golgi complex to the apical PM in polarized HA-MDCK cells treated with DMSO or 20 μM CRT0066101. The cells were subjected to HA transport assay as described in Methods, fixed, and visualized by mKate2 fluorescence and anti-GM130 immunostaining. The insets show magnified and merged images of mKate2-FM4-HA (magenta) and the Golgi marker GM130 (green) in the boxed areas. Scale bars, 5 µm (large panels) and 2.5 µm (insets). **(B)** Quantification of mKate2-FM4-HA transport from the Golgi complex to the apical PM. The ratio of fluorescence intensity of mKate2-FM4-HA in the apical PM and the Golgi complex, normalized as the values in DMSO-treated control (Cont) cells, as described in Methods, is shown. Boxes delimit the first and third quartiles, and the central line is the median, whereas the cross represents the mean value. The whiskers represent the minimum and maximum values after outlier removal (DMSO: n = 451 cells; CRT0066101: n = 378 cells; ****, P < 0.0001; unpaired two-tailed Student’s t test). **(C, D)** PAUF-MycHis secretion in polarized MDCK cells treated with DMSO or 20 μM CRT0066101. Polarized MDCK cells stably expressing PAUF-MycHis were pretreated with DMSO or 20 μM CRT0066101 at 37°C for 15 min and further incubated with fresh medium containing the respective reagent at 37°C for 3 h. The cell lysates and secreted proteins in the upper (apical) and lower (basolateral) compartments of the Transwell inserts were analyzed by immunoblotting (IB) with an anti-His antibody. **(D)** Quantification of apically secreted PAUF-MycHis. The amount of apically secreted PAUF-MycHis relative to the total cellular level and normalized as the values in DMSO-treated control cells is shown. Data are means ± SEM (n = 3 independent experiments; *, P = 0.192; unpaired two-tailed Student’s t test). Source data are available online for this figure.

We next investigated the impact of PKD inhibition on secretion of the CARTS cargo PAUF-MycHis in polarized MDCK cells. In control cells, the mature, higher-molecular-weight form of PAUF-MycHis was predominantly detected in the medium collected from the apical compartment (Fig. 4C). Treatment with CRT0066101 significantly decreased apical secretion of PAUF-MycHis to ∼36% of control levels (Fig. 4C,D). Altogether, these results indicate that PKD kinase activity is required not only for basolateral trafficking but also for efficient TGN-to-apical PM transport of selected cargoes, such as HA and PAUF, in polarized MDCK cells.

### Caveolins are required for the biogenesis of secretory carriers at the TGN

Our findings thus far indicate that the biogenesis of two distinct classes of TGN-derived transport carriers, CARTS and HA carriers, relies on lipid transfer at ER–Golgi MCSs and on PKD activity. However, how these factors coordinate cargo sorting, membrane budding and fission in the absence of conventional cytoplasmic coats remains unclear. We therefore turned to caveolin-1 (CAV1/VIP21), a cholesterol-binding integral membrane protein that was identified in HA carriers isolated from polarized MDCK cells (Kurzchalia *et al*, 1992; Glenney, 1992; Murata *et al*, 1995). Although early work implicated CAV1 in promoting TGN-to-apical PM transport of HA (Scheiffele *et al*, 1998), subsequent studies challenged this role (Manninen *et al*, 2005), leaving the function of CAV1 at the TGN unresolved.

To determine whether CAV1 is a component of HA carriers in our system, we employed a retention using selective hooks (RUSH) assay (Boncompain *et al*, 2012). As shown in Fig. EV4A, full-length CAV1a, the longer isoform of CAV1, was fused to a streptavidin (Str)-binding peptide (SBP) and EGFP. This chimera protein (CAV1-SBP-EGFP) was retained in the ER when co-expressed with the ER hook Str-Ii, and synchronously released from the ER upon addition of biotin or biotin methyl ester (BME; Uehara *et al*, 2025) (Fig. EV4B). At 20 min after transport initiation, CAV1-SBP-EGFP accumulated in the juxtanuclear Golgi area as well as in peripheral puncta, likely corresponding to TGN-derived transport carriers, before reaching the PM at later time points, where CAV1 forms caveolae. In HA-HeLa cells expressing CAV1-SBP-EGFP, ∼65% of HA carriers were positive for CAV1-SBP-EGFP (Fig. 5A,B), indicating that HA carriers incorporate CAV1. Co-expression of CAV1-SBP-EGFP with mKate2-FM4-PAUF similarly revealed CAV1 incorporation into CARTS, albeit to a slightly lesser extent (∼50%; Fig. 5A,B). Live-cell imaging after release from a 20°C block by incubation at 37°C showed that CAV1-SBP-EGFP and mKate2-FM4-PAUF were co-exported from the TGN (Movies EV2, EV3, and Appendix Fig. S2, arrowheads). This was frequently accompanied by the formation of CAV1-positive tubules. In relatively large CARTS, CAV1 could also be resolved covering the carrier surface, likely because their larger size allowed this organization to be spatially resolved by light microscopy (Movie EV3, and Appendix Fig. S2, arrows).

**Figure 5.**
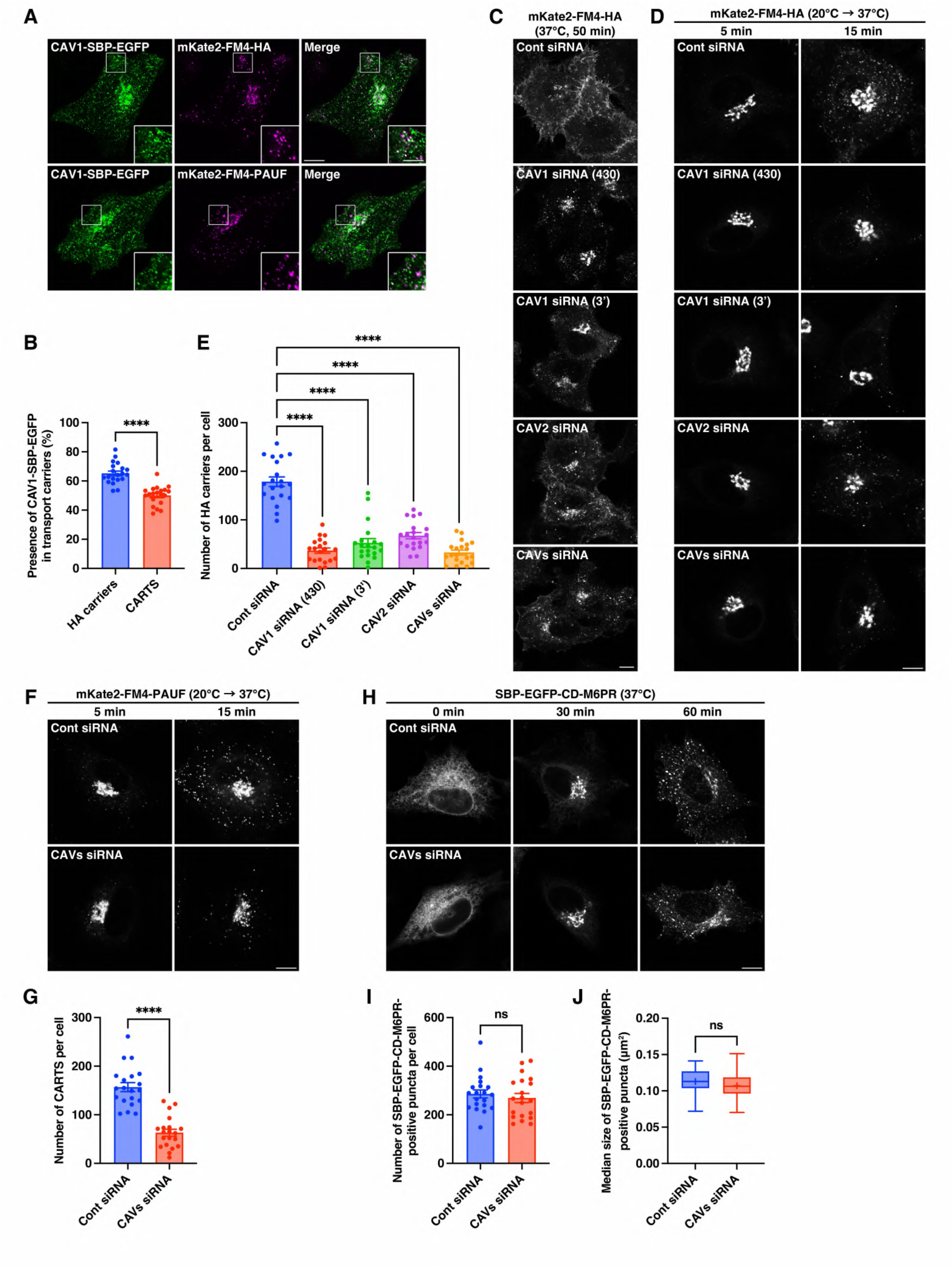
Cavs are required for the biogenesis of secretory carriers at the TGN. **(A, B)** Colocalization of CAV1-SBP-EGFP with HA carriers and CARTS. HA-HeLa and PAUF-HeLa cells expressing Str-Ii_CAV1-SBP-EGFP were subjected to cotransport assays as described in Methods, fixed, and visualized by EGFP and mKate2 fluorescence. The insets show magnified images of the boxed areas. See also Movies EV2 and EV3. **(B)** Quantification of the mKate2-FM4-HA–positive HA carriers containing CAV1-SBP-EGFP (blue), and the mKate2-FM4-PAUF–positive CARTS containing CAV1-SBP-EGFP (red). Data are means ± SEM, individual points represent values from individual cells ([HA carriers] mKate2-FM4-HA positive: n = 2,487 puncta and CAV1-SBP-EGFP positive: n = 22,939 puncta in 20 cells; [CARTS] mKate2-FM4-PAUF positive: n = 2,362 puncta and CAV1-SBP-EGFP positive: n = 26,523 puncta in 20 cells; ***, P < 0.0001; unpaired two-tailed Student’s t test). **(C)** mKate2-FM4-HA transport to the PM under control (Cont), CAV1, CAV2, and CAV1/2 (CAVs) knockdown conditions in HA-HeLa cells. The cells were incubated at 37°C for 50 min with D/D solubilizer and cycloheximide, fixed, and visualized by mKate2 fluorescence. **(D, E)** HA carrier biogenesis under control, CAV1, CAV2, and CAV1/2 knockdown conditions in HA-HeLa cells. The cells were subjected to HA carrier formation assay as described in Methods, fixed, and visualized by mKate2 fluorescence. **(E)** Quantification of the HA carrier biogenesis. The number of HA carriers at 15 min after the temperature shift to 37°C is shown. Data are means ± SEM (n = 20 cells per condition; ****, P < 0.0001; one-way ANOVA multiple comparison test). **(F, G)** CARTS biogenesis under control, CAV1, CAV2, and CAV1/2 knockdown conditions in PAUF-HeLa cells. The cells were subjected to CARTS formation assay as described in Methods, fixed, and visualized by mKate2 fluorescence. **(G)** Quantification of the CARTS biogenesis. The number of CARTS at 15 min after the temperature shift to 37°C is shown. Data are means ± SEM (n = 20 cells per condition; ****, P < 0.0001; unpaired two-tailed Student’s t test). **(H**–**J)** SBP-EGFP-CD-M6PR transport from the ER to endosomes via the Golgi complex under control and CAV1/2 knockdown conditions in HeLa cells expressing Str-KDEL_SBP-EGFP-CD-M6PR. The cells were subjected to CD-M6PR transport assay as described in Methods, fixed, and visualized by EGFP fluorescence. **(I, J)** Quantification of SBP-EGFP-CD-M6PR transport from the Golgi complex to endosomes. The number **(I)** and the median size **(J)** of SBP-EGFP-CD-M6PR–positive puncta at 30 min after transport initiation is shown. **(I)** Data are means ± SEM (n = 20 cells per condition; unpaired two-tailed Student’s t test). **(J)** Boxes delimit the first and third quartiles, and the central line is the median, whereas the cross represents the mean value. The whiskers represent the minimum and maximum values (n = 20 cells per condition; unpaired two-tailed Student’s t test). Scale bars, 10 µm (large panels) and 5 µm (insets). Source data are available online for this figure.

We next asked whether CAV1 is merely a passive cargo of HA carriers and CARTS or instead contributes to the molecular machinery driving their biogenesis. To address this, we examined the effects of siRNA-mediated depletion of the two CAV isoforms expressed in HeLa cells, CAV1 and CAV2. Two independent CAV1 siRNAs (430 and 3’) and one CAV2 siRNA efficiently reduced the corresponding protein levels, and combined depletion of CAV1 (siRNA 430) and CAV2 resulted in simultaneous loss of both proteins (Appendix Fig. S1B). CAV2, a regulator of CAV1 during caveolae formation (Sowa *et al*, 2003), was detected as a single prominent band in control cells, whereas a faint doublet band was observed in CAV1-depleted cells. This is consistent with previous reports showing that the CAV1–CAV2 hetero-oligomerization is required for stabilization and transport of CAV2 from the Golgi complex to the PM (Parolini *et al*, 1999; Mora *et al*, 1999). Under conditions of either single or combined knockdown of CAV1 and CAV2, synchronized TGN-to-PM transport of mKate2-FM4-HA was markedly delayed (Fig. 5C), and HA carrier biogenesis was significantly reduced (Fig. 5D,E). The relatively mild phenotype observed upon CAV2 depletion alone likely reflects its predominantly regulatory role. Combined CAV1/2 depletion, which caused the strongest inhibition of HA carrier biogenesis, similarly impaired CARTS biogenesis (Fig. 5F,G).

To assess the specificity of this effect and confirm that it does not affect general TGN function, we next tested whether CAV1/2 depletion affects clathrin-dependent TGN-to-endosome trafficking. For this purpose, we used the RUSH assay to monitor synchronized transport of the cation-dependent mannose 6-phosphate receptor (CD-M6PR) (Fig. 5H). Quantification of both the number and the median size of SBP-EGFP-CD-M6PR–positive puncta (corresponding to clathrin-coated vesicles and/or endosomal membranes) at 60 min revealed no significant differences between control and CAV1/2-depleted cells (Fig. 5I,J), indicating that clathrin-coated vesicle biogenesis at the TGN does not require CAVs. Together, these results identify CAVs as essential components of the machinery required for the biogenesis of secretory carriers at the TGN, while leaving clathrin-dependent TGN export unaffected.

### CAVs promote secretory carrier budding by oligomerizing and interacting with PKD

CAVs oligomerize progressively along the secretory pathway (Monier *et al*, 1995). Current models propose that CAV monomers or small oligomers exit the ER and assemble into 8S complexes (∼11–14 protomers) upon arrival at the Golgi membranes (Morales-Paytuví *et al*, 2023). The 8S complexes further assemble into 70S complexes (∼150 protomers) in a cholesterol-dependent manner before export from the TGN to the PM (Hayer *et al*, 2010). At the PM, cavin proteins are recruited to 70S CAV complexes to form stable caveolae (Hayer *et al*, 2010; Parton, 2018). Although cavins are essential for caveolae formation in vivo (Hill *et al*, 2008; Liu *et al*, 2008), expression of CAV1 alone in a bacterial model system is sufficient to induce membrane curvature and coated vesicle budding (Walser *et al*, 2012). Recent computational and experimental studies further suggest that the CAV1-8S complex generates membrane curvature by asymmetric insertion into one membrane leaflet, displacing lipids from the opposing leaflet and inducing local lipid tilting and bending, thereby driving cholesterol concentration and lipid sorting in surrounding membrane (Doktorova *et al*, 2025; Barnoy *et al*, 2025). Negatively curved lipids such as cholesterol and DAG are predicted to enhance this process, underscoring their mechanical contribution to CAV1-driven membrane deformation (Barnoy *et al*, 2025).

Based on these observations, we hypothesized that CAV oligomers act as a coat-like scaffold at the TGN together with lipid nanodomains to promote secretory carrier budding. To test this hypothesis, we used the dominant-negative CAV1 P132L mutant, which disrupts protomer-protomer interactions and prevents proper CAV oligomerization, resulting in its accumulation in the Golgi complex (Lee *et al*, 2002; Han *et al*, 2023). In HA-HeLa cells expressing CAV1 WT-mEGFP, synchronized mKate2-FM4-HA trafficking proceeded normally, with HA reaching the PM by 50 min (Fig. 6A,B). By contrast, expression of CAV1 P132L-mEGFP caused pronounced retention of mKate2-FM4-HA in the juxtanuclear Golgi region, where the two proteins showed strong colocalization. Moreover, CAV1 P132L-mEGFP expression inhibited the formation of GST-PKD2 KD-induced membrane tubules (Fig. 6C,D). As these tubules represent stabilized budding membranes due to a fission defect caused by PKD2 KD expression, our results suggest that the capacity of CAV1 to form oligomers is required for the membrane budding process at the TGN. Of note, these phenotypes were not observed upon expression of the dominant-negative CAV1 Y14F mutant (Fig. 6A–D). Since this mutant inhibits caveolae-dependent endocytosis by preventing the Tyr14 phosphorylation-dependent conformational rearrangement of the caveolar coat required for caveolar swelling and release from the plasma membrane (Zimnicka *et al*, 2016), our results argue against the possibility that the effects of CAV1 P132L are an indirect consequence of impaired caveolae-dependent endocytosis or recycling.

**Figure 6.**
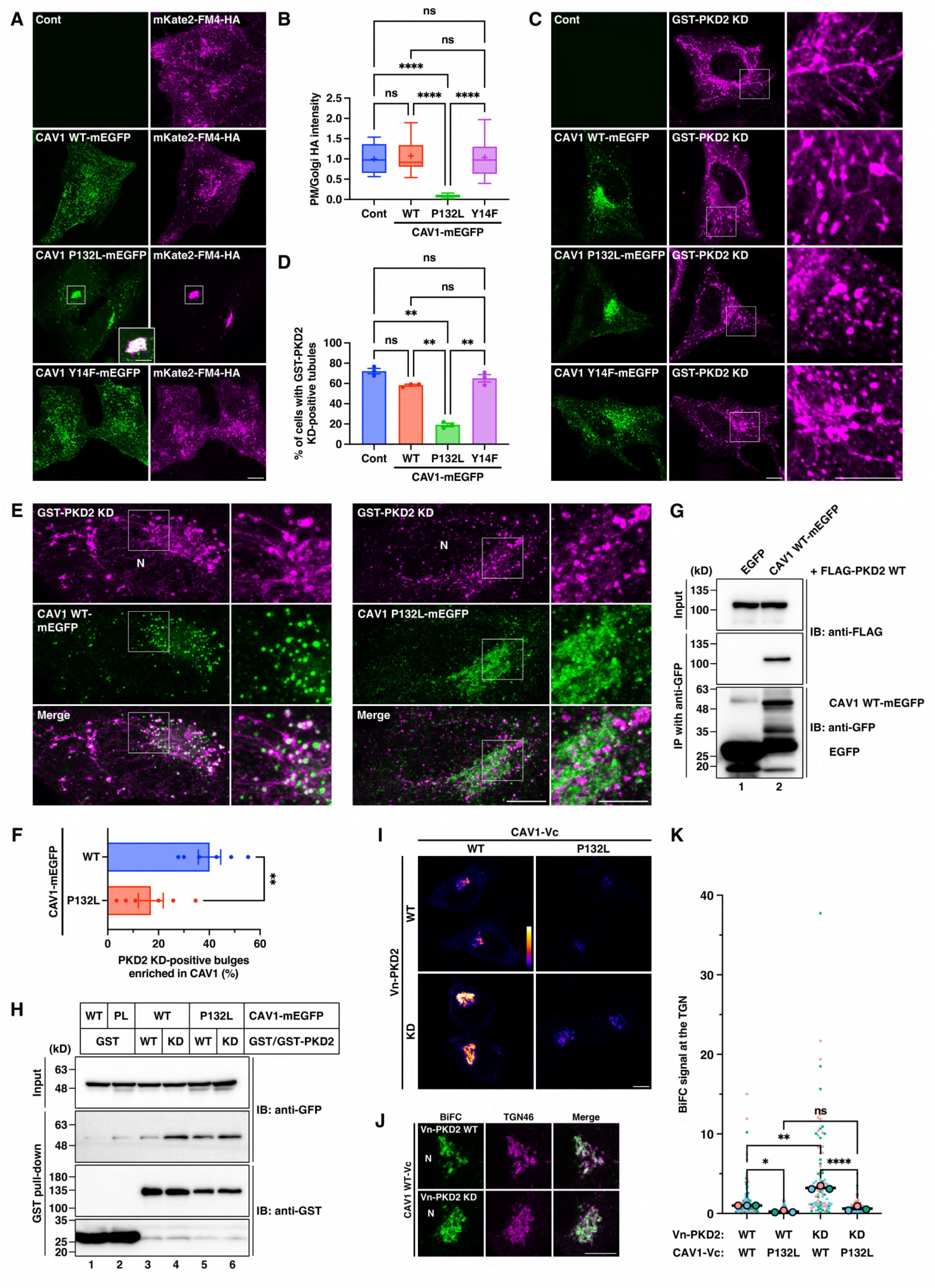
CAVs promote secretory carrier budding by oligomerizing and interacting with PKD. **(A, B)** Accumulation of mKate2-FM4-HA at the Golgi complex due to CAV1 P132L-mEGFP expression. Control (Cont) HA-HeLa cells and HA-HeLa cells expressing each CAV1-mEGFP construct were incubated at 37°C for 50 min with D/D solubilizer and cycloheximide, fixed, and visualized by mEGFP and mKate2 fluorescence. The inset shows a magnified and merged image of CAV1 P132L-mEGFP and mKate2-FM4-HA in the boxed area. Scale bars, 10 µm (large panels) and 5 µm (inset). **(B)** Quantification of mKate2-FM4-HA transport from the Golgi complex to the PM. The ratio of fluorescence intensity of mKate2-FM4-HA in the PM and the Golgi complex, normalized as the values in control cells, as described in Methods, is shown. Boxes delimit the first and third quartiles, and the central line is the median, whereas the cross represents the mean value. The whiskers represent the minimum and maximum values after outlier removal (n = 12 cells per condition; ****, P < 0.0001; one-way ANOVA multiple comparison test). **(C, D)** Membrane tubule formation in HeLa cells expressing GST-PKD2 KD alone (control) or co-expressing GST-PKD2 KD and CAV1 (WT, P132L, or Y14F)-mEGFP. The cells were fixed and visualized by GFP fluorescence and anti-GST immunostaining. Single-plane images are shown with magnified views of the boxed areas (right column). Scale bars, 10 µm. **(D)** Quantification of the GST-PKD2 KD-induced membrane tubule formation. The percentage of cells with GST-PKD2 KD-positive tubules is shown. Data are means ± SEM (n = 3 independent experiments [100–108 cells per condition]; **, control vs. P132L, P = 0.0017; WT vs. P132L, P = 0.0011; P132L vs. Y14F, P = 0.0054; one-way ANOVA multiple comparison test). **(E, F)** Accumulation of CAV1 WT-mEGFP at bulging TGN domains connected to GST-PKD2 KD-positive tubules. HeLa cells co-expressing GST-PKD2 KD with CAV1 (WT or P132L)-mEGFP were fixed and visualized by GFP fluorescence and anti-GST immunostaining. 3D super-resolution microscopic images were acquired as described in Methods. Maximum intensity merges of Z-stack images are shown with magnified views of the boxed areas (right of each panel). Arrowheads indicate CAV1 WT-mEGFP–positive bulging TGN domains connected to GST-PKD2 KD-positive tubules. N, nucleus. Scale bars, 10 µm (left panels) and 5 µm (right panels). See also Movie EV4. **(F)** Quantification of CAV1 accumulation at the bulging TGN domains. The percentage of GST-PKD2 KD-positive bulges enriched in CAV1 (WT or P132L)-mEGFP is shown. Histograms of the measured intensities were generated (Appendix Fig. S3A,B), and PKD2 KD-positive bulges highly enriched in CAV1 signal (as defined as relative enrichment >5) were classified as CAV1-enriched. Data are means ± SEM ([CAV1 WT] PKD2 KD positive: n = 47–74 bulges in 6 cells; [CAV1 P132L] PKD2 KD positive: n = 15–58 bulges in 6 cells; **, P = 0.0056; unpaired two-tailed Student’s t test). **(G)** Interaction of FLAG-PKD2 WT with CAV1 WT-mEGFP. HEK 293T cells stably expressing FLAG-PKD2 WT were transfected with a plasmid for EGFP (control) or CAV1 WT-mEGFP. After 24 h, the cells were subjected to immunoprecipitation (IP) with an anti-GFP nanobody as described in Methods. The cell lysates (Input) and precipitates were immunoblotted (IB) with the indicated antibodies. **(H)** Interaction of CAV1 (WT or P132L)-mEGFP with GST-PKD2 (WT or KD). HEK 293T cells co-expressing GST (control) or GST-PKD2 (WT or KD) with CAV1 (WT or P132L [PL])-mEGFP were subjected to GST pull-down as described in Methods. The cell lysates (Input) and precipitates were immunoblotted with the indicated antibodies. (**I**–**K**) BiFC visualization of the CAV1-PKD2 interaction at the TGN. HeLa Tet-On cells co-expressing Vn-PKD2 (WT or P132L) with CAV1 (WT or P132L)-Vc were subjected to BiFC analysis as described in Methods. **(I)** Intensity of the BiFC signal. **(J)** Colocalization of the BiFC signal with TGN46. N, nucleus. Scale bars, 10 µm. **(K)** Quantification of the BiFC signal at the TGN. SuperPlot showing the indicated cell measurements (small symbols; 22–37 cells per condition for each biological replicate) and the median value for each independent biological replicate (larger, black-outlined circles; n = 3). Each color represents a different experimental replicate. A repeated measures one-way ANOVA test was performed using Tukey’s post-hoc multiple comparison test (*, P = 0.0262; **, P = 0.0094; ****, P < 0.0001). Source data are available online for this figure.

Three-dimensional super-resolution microscopy further revealed that CAV1 WT, but not the P132L mutant, accumulates at bulging TGN domains from which PKD2 KD-positive tubules emerge (Fig. 6E, arrowheads; 6F; Appendix Fig. S3; Movie EV4). By contrast, neither mKate2-FM4-HA (Fig. EV5A) nor TGN46 (Fig. EV5B) displayed comparable localized enrichment at these sites. Considering the previous finding that similar bulging domains are observed upon the halt of TGN export by a 20°C block (Ladinsky *et al*, 2002), our results suggest that CAV1 oligomers specifically concentrate at TGN exit domains amplified by the expression of PKD2 KD.

Finally, we examined whether CAV1 interacts with PKD2, potentially enabling their coordinated action during secretory carrier biogenesis at the TGN. Immunoprecipitation of expressed EGFP or CAV1 WT-mEGFP from HEK 293T cells stably expressing FLAG-PKD2 WT showed a specific association between PKD2 WT and CAV1 WT (Fig. 6G). Furthermore, GST pull-down assays using lysates of HEK 293T cells transiently expressing CAV1 (WT or P132L)-mEGFP together with GST-PKD2 (WT or KD) revealed that both WT and P132L CAV1-mEGFP were coprecipitated with GST-PKD2 KD and, to a lesser extent, PKD2 WT (Fig. 6H, lanes 3–6). The stronger association observed with PKD2 KD than with PKD2 WT may reflect stabilization of an otherwise transient interaction upon arrest of secretory carrier biogenesis. Because detergent solubilization may disrupt CAV1 oligomerization and oligomer-dependent interactions, we next examined CAV1–PKD2 interaction in situ by using bimolecular fluorescence complementation (BiFC). A BiFC signal generated by the interaction between Vn (N-terminal fragment of Venus)-PKD2 WT and CAV1 WT-Vc (C-terminal fragment of Venus) was detected at the TGN and was significantly enhanced by PKD2 KD expression (Fig. 6I, left column; 6J,K), consistent with the GST pull-down results (Fig. 6H). Notably, the BiFC signal was significantly reduced when CAV1 P132L-Vc was used (Fig. 6I, right column; 6K). Given the stronger localization of this mutant to the Golgi complex compared with CAV1 WT (Fig. 6A), our results indicate that Golgi localization and spatial proximity alone are not sufficient to generate a strong BiFC signal. This finding supports the specificity of the CAV1-PKD2 interactions and suggests that CAV1 oligomerization promotes interaction with PKD2 at the TGN.

Collectively, our results support a model in which CAV oligomerization establishes a curvature-inducing, membrane-embedded scaffold at lipid nanodomain-enriched TGN regions. Together with its interaction with PKD2, this scaffold facilitates coordinated cargo sorting, membrane budding and fission during secretory carrier biogenesis at the TGN.

## Discussion

The discovery of three classes of cytoplasmic coats –clathrin (Pearse, 1975), COPI (Malhotra *et al*, 1989), and COPII (Barlowe *et al*, 1994)– transformed our understanding of membrane trafficking. By contrast, the coat architecture of TGN-to-cell surface carriers has remained unclear, representing a long-standing obstacle to mechanistic analysis of this pathway. Notably, early high-voltage electron microscopy and 3D tomography studies in NRK cells by Howell and colleagues (Ladinsky *et al*, 1994; Wang *et al*, 1995) revealed a lace-like coat, distinct from the clathrin coat, that appeared to associate with secretory carrier biogenesis at the TGN. Although the molecular components of this structure were not identified, subsequent high-resolution 3D tomography studies from the same group (Ladinsky *et al*, 1999, 2002) showed that in NRK cells, the two trans-most Golgi cisternae, one containing lace-like coats and the other clathrin coats, form close contacts with the ER membrane. During a 20°C block, the ER was found to closely wraps around large bulging domains of the TGN. These bulging domains are thought to represent TGN exit sites, which become enlarged as a consequence of arrested TGN export. The contacts described in these studies are now recognized as ER–Golgi MCSs, which mediate non-vesicular lipid transfer to control Golgi lipid homeostasis and the biogenesis of secretory carriers, including CARTS (Goto *et al*, 2020; Venditti *et al*, 2020; Wakana & Campelo, 2021; Subra *et al*, 2023).

In the present study, we identify novel CARTS-like carriers transporting the apical PM cargo HA. At present, beyond HA transport, the broader functions and physiological relevance of these carriers remain to be determined. They may represent a constitutively active TGN export pathway that accommodates HA upon its exogenous expression. Alternatively, we cannot exclude the possibility that expression of HA itself enhances or induces a carrier population different from the CARTS pool. However, here we reveal a previously unrecognized mechanism of secretory carrier biogenesis at the TGN. Based on our data, we propose a model in which a PKD–CAV axis coordinates cargo sorting, membrane budding and fission downstream of lipid transfer at ER–Golgi MCSs (Fig. 7). This model comprises the following steps. (1) Lipid transfer at ER–Golgi MCSs locally increases cholesterol, SM, and DAG levels at the trans-Golgi/TGN regions closely apposed to the ER. Concomitantly, the SCAP– SREBP complex regulates cholesterol/PI4P exchange in an ER cholesterol-dependent manner via interaction with the VAP–OSBP–Sac1 complex (Wakana *et al*, 2021). (2) Cholesterol and SM assemble into lipid nanodomains while DAG recruits PKD to the TGN. (3) These lipid nanodomain-enriched TGN regions serve as molecular platforms for the functional assembly of CAV-8S complexes, selective secretory cargoes and/or cargo receptors, and the machineries required for cargo processing and sorting. Our observation that PAUF and its cargo receptor TGN46 (Lujan *et al*, 2024), but not HA, are incorporated into TGN-derived membrane tubules upon PKD inhibition (Figs. 3F–H,J,K, EV1A, and EV5) implicates PKD in selective cargo sorting and incorporation into nascent secretory carriers. (4) CAV-8S complexes further assemble into larger oligomeric structures, which may drive membrane budding through interactions with lipid nanodomains and/or with PKD. Consistent with this, perturbation of CAV oligomerization by the CAV1 P132L mutant –which does not interact with PKD2 in intact cells (Fig. 6I–K)– suppresses TGN-to-PM transport of HA (Fig. 6A,B) and the formation of PKD2 KD-induced tubules that correspond to secretory carrier precursors (Fig. 6C,D). Finally, (5) PKD activity promotes membrane fission, leading to (6) release of secretory carriers into the cytoplasm and completion of carrier biogenesis.

**Figure 7.**
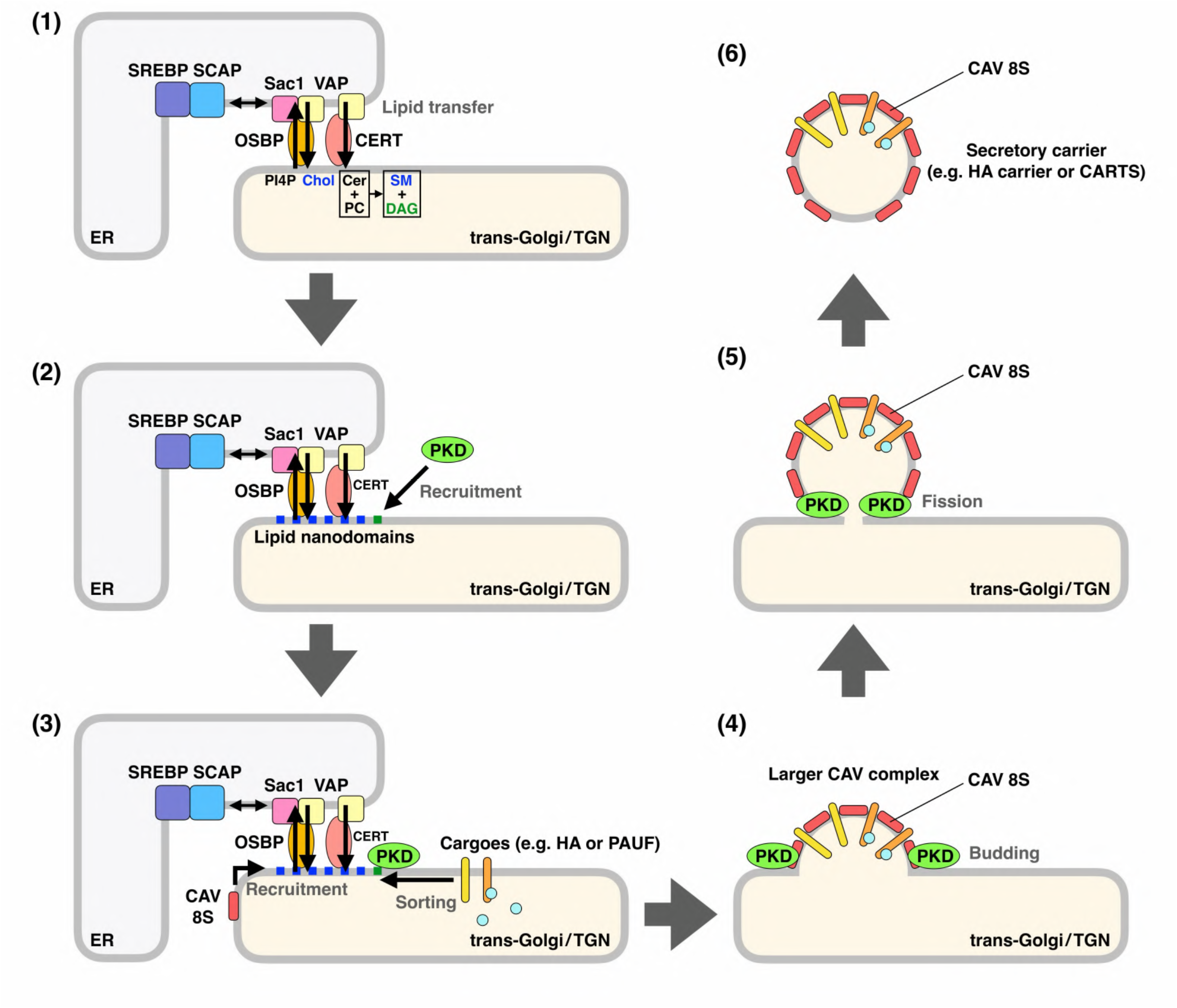
Working model for ER contact-induced secretory carrier biogenesis at the TGN. **(1)** Lipid transfer at ER–Golgi MCSs and a local increase in cholesterol (Chol), SM, and DAG levels at the trans-Golgi/TGN. **(2)** Assembly of cholesterol and SM into lipid nanodomains and DAG-dependent recruitment of PKD to the TGN. **(3)** Functional assembly of CAV-8S complexes, selective secretory cargoes (e.g. HA or PAUF) and/or cargo receptors, and the machineries required for cargo processing and sorting (not illustrated) at lipid nanodomain-enriched TGN regions. **(4)** Further assembly of CAV-8S complexes into larger oligomeric structures, which can drive membrane budding through interactions with lipid nanodomains and/or with PKD. **(5)** PKD-mediated membrane fission. **(6)** Secretory carrier release into the cytoplasm. Cer, ceramide; PC, phosphatidylcholine.

The relevance of ER–Golgi MCSs in secretory carrier biogenesis is supported by both lipid transfer and structural evidence. Both knockdown of ER–Golgi MCS components and OSW-1–mediated inhibition of OSBP-dependent cholesterol/PI4P exchange impaired the biogenesis of HA carriers (Fig. 2D–G) and CARTS (Fig. EV1) (Wakana *et al*, 2015, 2021). So far, all lipid transfer events at ER–Golgi MCSs appear to contribute to TGN export to varying degrees. Previous studies from our group and others have shown that the sensitivity to cholesterol and PI4P levels differs depending on the cargo (Venditti *et al*, 2019; Wakana *et al*, 2021; Kovács *et al*, 2023). Notably, perturbation of the sphingolipid-glycerolipid balance was recently shown to alter TGN membrane order and impair CAV1 trafficking to the plasma membrane (Carpentier *et al*, 2025). Moreover, our finding that expression of OSBP PH-FFAT, which stabilizes ER–Golgi MCSs while blocking OSBP-dependent cholesterol transport (Mesmin *et al*, 2013), markedly inhibits the export of HA from the TGN (Fig. 2H,I) and CARTS biogenesis (Wakana *et al*, 2015), suggests that the dynamic assembly and disassembly of ER–Golgi MCSs are necessary for the biogenesis of both types of secretory carriers. This notion is further supported by our previous finding that CARTS form at sites immediately adjacent to the VAP-A–OSBP complex-containing ER–Golgi MCSs (Wakana *et al*, 2021). Collectively, these observations suggest that efficient carrier formation depends not only on lipid transfer itself, but also on the dynamic organization of ER–Golgi MCSs, with the contribution of individual lipids possibly varying according to the cargo.

CAVs localize not only to the PM, where they scaffold caveolae, but also to the Golgi complex (Rothberg *et al*, 1992; Dupree *et al*, 1993; Mora *et al*, 1999). Earlier work by Ikonen and colleagues showed that anti-CAV1 antibodies inhibit TGN-to-apical PM transport of HA in permeabilized polarized MDCK cells (Scheiffele *et al*, 1998), suggesting a direct role for CAV1 in TGN export. This view was supported by the observation that glycosylphosphatidylinositol (GPI)-anchored proteins are preferentially retained in the Golgi complex in CAV1-null mouse embryo fibroblasts (Sotgia *et al*, 2002). However, subsequent work by Simons and colleagues showed that stable CAV1 knockdown does not inhibit HA or GPI-anchored protein transport from the TGN to the apical PM (Manninen *et al*, 2005), challenging a universal requirement for CAV1 in TGN export. Nevertheless, the delivery of other cargoes to the PM, including dysferlin (Hernández-Deviez *et al*, 2006, 2008), the angiotensin receptor (Wyse *et al*, 2003), the insulin receptor (Cohen *et al*, 2003), and the stretch-activated channel TRPC1 (Brazer *et al*, 2003), appears to depend on CAVs (Parton & Simons, 2007). Importantly, export of dysferlin from the TGN has been suggested to occur via both CAV-dependent and independent routes (Hernández-Deviez *et al*, 2008). Such context dependence may apply to PAUF as well: in our study, depletion of CAV1/2 produced relatively modest defects in CARTS biogenesis (Fig. 5F,G), compared with the more pronounced inhibition of HA carrier formation (Fig. 5D,E). This is consistent with the slightly lower incorporation efficiency of CAV1 into CARTS compared with HA carriers (Fig. 5A,B). Functional redundancy among parallel secretory routes may therefore partially mask the contribution of CAVs and complicate assessment of their role at the TGN.

Our results also revealed a previously unappreciated role for PKD2 in selective cargo sorting and CAV-mediated membrane budding, complementing its established function in membrane fission and supporting the view that PKD acts as a central regulator of secretory carrier biogenesis at the TGN (Wakana & Campelo, 2021). PKD phosphorylates and activates phosphatidylinositol 4-kinase IIIβ to produce PI4P (Hausser *et al*, 2005), which recruits pleckstrin homology (PH)-domain containing proteins such as CERT and OSBP to the trans-Golgi/TGN. Subsequent PKD-mediated phosphorylation of CERT and OSBP promotes their dissociation from the Golgi membranes (Fugmann *et al*, 2007; Nhek *et al*, 2010), likely enabling the dynamic assembly and disassembly of ER–Golgi MCSs at sites of secretory carrier formation. In addition to its established role in basolateral trafficking (Yeaman *et al*, 2004), our data demonstrate that PKD kinase activity is also required for efficient apical transport (Fig. 4). PKD has also been implicated in regulated secretion of insulin (Sumara *et al*, 2009) and chromogranin A (Cruz-Garcia *et al*, 2013), as well as in mechanically adaptive regulation of constitutive Golgi export (Bhaskar Naidu *et al*, 2026), raising the possibility that the PKD–CAV axis described here may operate more broadly across distinct modes of regulated secretory traffic.

In conclusion, our findings uncover an unexpected functional connection between PKD and CAV at the TGN and define a lipid-driven, membrane-embedded scaffold mechanism that supports coatless carrier formation. This work provides a conceptual framework for understanding how lipid nanodomains and integral membrane proteins can substitute for classical cytoplasmic coats and opens up new directions for dissecting the molecular basis of secretory carrier formation.

## Methods

### Reagent and tools table

| Reagent/Resource | Reference or Source | Identifier or Catalog Number |
|---|---|---|
| **Experimental Models** |  |  |
| HeLa cells (*Homo sapiens*) | ATCC | CCL-2 |
| MDCK cells (*Canis lupus familiaris*) | ATCC | CCL-34 |
| HEK 293T cells (*Homo sapiens*) | Gift from S. Hirose (Tokyo Institute of Technology, Japan) |  |
| PLAT-A packaging cells | Kitamura *et al*, 2003 |  |
| HA-HeLa cells | This study | “Methods” |
| HA-MDCK cells | This study | “Methods” |
| PAUF-HeLa cells | Wakana & Tagaya, 2023 |  |
| PAUF-MycHis-HeLa cells | This study | “Methods” |
| PAUF-MycHis-MDCK cells | This study | “Methods” |
| FLAG-PKD2 WT-HEK 293T cells | This study | “Methods” |
| HeLa Tet-On cells | This study | “Methods” |
| HeLa Tet-On HA-mEGFP cells | This study | “Methods” |
| **Recombinant DNA** |  |  |
| Influenza-FPV-HA-mEGFP | Addgene | Gift from S. Chiantia (#127810; Dunsing et al., 2018 |
| CD-MPR-GFP-RUSH (Str-KDEL_IRES_SBP-EGFP-CD-MPR) | Addgene | Gift from J. Bonifacino (#202797; Chen et al., 2017) |
| Str-Ii_SBP-EGFP-Golgin84 | Addgene | Gift from F. Perez (#65303; Boncompain et al., 2012) |
| Str-Ii_CAV1-SBP-EGFP | This study | “Methods” |
| CAV1 WT-mEGFP | Addgene | Gift from A. Helenius (#27704; Hayer et al., 2010) |
| CAV1 P132L-mEGFP | Addgene | Gift from A. Helenius (#27708; Hayer et al., 2010) |
| CAV1 Y14F-mEGFP | This study | “Methods” |
| Rab6a-GFP | Gift from F. A. Barr (University of Oxford, Oxford, UK) |  |
| Rab8a-GFP | Gift from F. A. Barr (University of Oxford, Oxford, UK) |  |
| Rab11a-GFP | Gift from F. A. Barr (University of Oxford, Oxford, UK) |  |
| EQ-SM (tagged with oxGFP) | Gift from C.G. Burd (Yale School of Medicine, New Haven, CT) (Deng *et al*, 2016) |  |
| Myc-OSBP PH-FFAT | (Wakana *et al*, 2015) |  |
| GST | Gift from V. Malhotra (Centre for Genomic Regulation, Barcelona, Spain) |  |
| GST-PKD2 WT | Gift from V. Malhotra (Centre for Genomic Regulation, Barcelona, Spain) |  |
| GST-PKD2 KD (K580N) | Gift from V. Malhotra (Centre for Genomic Regulation, Barcelona, Spain) |  |
| pcDNA3.1(+)-PAUF-MycHis | Gift from S.S. Koh (Korea Research Institute of Bioscience and Biotechnology, Daejeon, Korea) (Kim *et al*, 2009) |  |
| pCI-IRES-Bsr | Yonekawa *et al*, 2011 |  |
| pCI-PAUF-MycHis-IRES-Bsr | This study | “Methods” |
| pCX4-IRES-Bsr | Akagi *et al*, 2003 |  |
| pCX4-ss-mKate2-FM4-HA-IRES-Bsr | This study | “Methods” |
| pCX4-ss-mKate2-FM4-PAUF-IRES-Bsr | Wakana & Tagaya, 2023 |  |
| pCX4-FLAG-PKD2-IRES-Bsr | This study | “Methods” |
| pMXs-IRES-Puro | Cell Biolabs | RTV-014 |
| pMXs-PAUF-MycHis-IRES-Puro | This study | “Methods” |
| pRetroX-Tet-On Advanced | Clontech | Retro-X Tet-On Advanced Inducible Expression System (632104) |
| pRetroX-Tight-Pur | Clontech | Retro-X Tet-On Advanced Inducible Expression System (632104) |
| pRetroX-Tight-HA-mEGFP-Pur | This study | “Methods” |
| pCX4-Vn-PKD2 (WT, KD)-IRES-Bsr | This study | “Methods” |
| pRetroX-Tight-CAV1 (WT, P132L)-Vc-Pur | This study | “Methods” |
| **Antibodies** |  |  |
| FLAG (mouse monoclonal) | Sigma-Aldrich | F3165 |
| FLAG (rabbit polyclonal) | Sigma-Aldrich | F7425 |
| tRFP (mKate2) (rabbit polyclonal) | Evrogen | AB233 |
| Myc (mouse monoclonal) | Santa Cruz Biotechnology | sc-40 |
| GFP (mouse monoclonal) | Santa Cruz Biotechnology | sc-9996 |
| GFP (Vn) (rabbit monoclonal) | Cell Signaling | 2956 |
| GFP (Vc) (mouse monoclonal) | FUJIFILM Wako Pure Chemical Corporation | 012-22541 |
| GST (rabbit polyclonal) | Santa Cruz Biotechnology | sc-459 |
| Penta-His (mouse monoclonal) | Qiagen | 34660 |
| Calnexin (mouse monoclonal) | BD Biosciences | 610524 |
| TGN46 (rabbit polyclonal) | Wakana *et al*, 2021 |  |
| TGN46 (sheep polyclonal) | Bio-Rad | AHP500GT |
| GM130 (mouse monoclonal) | BD Biosciences | 610823 |
| Clathrin HC (goat polyclonal) | Santa Cruz Biotechnology | sc-6579 |
| γ-adaptin (mouse monoclonal) | BD Biosciences | 610385 |
| β-COP (rabbit polyclonal) | Gift from H.-P. Hauri (University of Basel, Basel, Switzerland) |  |
| Sec31 (rabbit polyclonal) | Hirose *et al*, 2004 |  |
| PKD2 (rabbit monoclonal) | Cell Signaling | 8188 |
| PKD3/PKC nu (rabbit monoclonal) | Cell Signaling | 5655 |
| CAV1 (rabbit polyclonal) | Proteintech | 16447-1-AP |
| CAV2 (rabbit polyclonal) | GeneTex | GTX108294 |
| α-tubulin (mouse monoclonal) | Sigma-Aldrich | T6074 |
| ZO-1 (rabbit polyclonal) | Proteintech | 21773-1-AP |
| Goat anti-rabbit IgG, Alexa Fluor 350 | Thermo Fisher Scientific | A11046 |
| Donkey anti-rabbit IgG, Alexa Fluor 350 | Thermo Fisher Scientific | A10039 |
| Goat anti-mouse IgG, Alexa Fluor 488 | Thermo Fisher Scientific | A11029 |
| Goat anti-rabbit IgG, Alexa Fluor 488 | Thermo Fisher Scientific | A11034 |
| Donkey anti-rabbit IgG, Alexa Fluor 488 | Thermo Fisher Scientific | A21206 |
| Donkey anti-goat IgG, Alexa Fluor 488 | Thermo Fisher Scientific | A11055 |
| Goat anti-mouse IgG, Alexa Fluor 594 | Thermo Fisher Scientific | A11032 |
| Goat anti-rabbit IgG, Alexa Fluor 594 | Thermo Fisher Scientific | A11037 |
| Donkey anti-sheep IgG, Alexa Fluor 594 | Thermo Fisher Scientific | A11016 |
| Goat anti-mouse IgG, Alexa Fluor 647 | Thermo Fisher Scientific | A21235 |
| Goat anti-rabbit IgG, Alexa Fluor 647 | Thermo Fisher Scientific | A21245 |
| Goat anti-mouse IgG, HRP | Bio-Rad | 170-6516 |
| Goat anti-rabbit IgG, HRP | Bio-Rad | 170-6515 |
| **Oligonucleotides and other sequence-based reagents** |  |  |
| See Appendix Table S1 |  |  |
| **Chemicals, Enzymes and other reagents** |  |  |
| D/D solubilizer | Clontech | 635054 |
| Biotin | Sigma-Aldrich | B4501 |
| BME | MedChemExpress | HY-W107202 |
| Cycloheximide | FUJIFILM Wako Pure Chemical Corporation | 037-20991 |
| DMSO | FUJIFILM Wako Pure Chemical Corporation | 046-21981 |
| OSW-1 | Gift from Y. Mimaki (Tokyo University of Pharmacy and Life Sciences, Tokyo, Japan) |  |
| CRT0066101 | Cayman Chemical | 15337 |
| OPTI-MEM | Thermo Fisher Scientific | 31985-070 |
| DMEM | FUJIFILM Wako Pure Chemical Corporation | 043-30085 |
| FBS | Sigma-Aldrich | 173012 |
| Blasticidin S | Kaken Pharmaceutical | KK-400 |
| Puromycin | FUJIFILM Wako Pure Chemical Corporation | 160-23151 |
| G418 | FUJIFILM Wako Pure Chemical Corporation | 074-05963 |
| Doxycycline | Sigma-Aldrich | D-9891 |
| X-tremeGENE 9 DNA transfection reagent | Roche | 6365787001 |
| Oligofectamine transfection reagent | Thermo Fisher Scientific | 12252011 |
| Lipofectamine 2000 transfection reagent | Thermo Fisher Scientific | 11668500 |
| Polyethylenimine | Polysciences | 24765 |
| Benzonase nuclease | Sigma-Aldrich | E8263 |
| GFP-Trap Agarose | ChromoTek | gta |
| Glutathione Sepharose 4B | Cytiva | 17075601 |
| e-PAGEL HR 5-20% | ATTO | EHR-T520L |
| WSE-7210 EzFastBlot HMW | ATTO | 2332595 |
| Immobilon Classico Western HRP substrate | Millipore | WBLUC0500 |
| Signal Enhancer HIKARI for Western Blotting and ELISA | Nacalai Tesque | 02267-41 |
| **Software** |  |  |
| ImageJ | National Institutes of Health |  |
| Fiji | Schindelin *et al*, 2012 |  |
| MATLAB R2024b | MathWorks http://www.mathworks.com/products/matlab/ | RRID:SCR_001622 |
| GraphPad Prism (v. 11.1.0) | GraphPad |  |
| FV10-ASW | Olympus |  |
| FV31S-SW | Olympus |  |
| cellSens FV | Evident |  |
| cellSens Dimension | Evident |  |
| Microsoft Excel (v.16.112.4) | Microsoft |  |
| **Other** |  |  |
| Fluoview FV1200 confocal microscope | Olympus |  |
| Fluoview FV3000 confocal microscope | Olympus |  |
| Fluoview FV4000 confocal microscope | Evident |  |
| IXplore IX85 Spin confocal spinning disk microscope | Evident |  |

### Plasmid construction

For establishment of HA-HeLa and HA-MDCK cells, the cDNAs encoding the signal sequence (ss) of human growth hormone 1 (GH1) (aa 1-26)-mKate2-FM4 and influenza-FPV-HA (aa 491-563) were inserted into pCX4-IRES-Bsr. For establishment of HeLa and MDCK cells stably expressing PAUF-MycHis, the cDNA encoding PAUF-MycHis was inserted into pCI-IRES-Bsr and pMXs-IRES-Puro, respectively. For establishment of a HeLa Tet-On HA-mEGFP cell line, the cDNA encoding HA-mEGFP was inserted into pRetroX-Tight-Pur. For establishment of HEK 293T cells stably expressing FLAG-PKD2 WT, the cDNAs encoding FLAG-PKD2 WT was inserted into pCX4-IRES-Bsr. For co-expression of CAV1-SBP-EGFP and Str-Ii, the cDNA encoding Golgin84 was removed from the plasmid for Str-Ii_SBP-EGFP-Golgin84 and the cDNA encoding CAV1 was inserted into the plasmid upstream of SBP-EGFP coding region. For expression of CAV1 Y14F-mEGFP, a point mutation was introduced into CAV1 WT-mEGFP by PCR using primers designed for replacing tyrosine 14 with phenylalanine. For BiFC analysis, the cDNAs encoding Vn (aa 1–172 of Venus) and human PKD2 (WT or KD [K580N]) were inserted into pCX4-IRES-Bsr, whereas the cDNAs encoding CAV1 (WT or P132L) and Vc (aa 154–238 of Venus) were inserted into pRetroX-Tight-Pur.

### Cell culture and transfection

HeLa, MDCK, and HEK 293T cells were grown in DMEM supplemented with 10% FBS. PLAT-A packaging cells were grown in DMEM supplemented with 10% FBS, 10 μg/ml blasticidin S, and 1 μg/ml puromycin. Plasmid and siRNA transfection into HeLa cells was performed using X-tremeGENE 9 DNA transfection reagent and Oligofectamine transfection reagent, respectively, according to the manufacturers’ protocols. Plasmid transfection into HEK 293T and PLAT-A packaging cells was performed using Lipofectamine 2000 transfection reagent and polyethylenimine, respectively.

### Establishment of stable cell lines

For establishment of HA-HeLa and HA-MDCK cells, PLAT-A packaging cells were transfected with pCX4-ss-mKate2-FM4-HA-IRES-Bsr, and 48 h later, the medium containing retrovirus was collected and used for infection of HeLa and MDCK cells, respectively. Selection of the stable cell lines was performed with 10 μg/ml blasticidin S. A HEK 293T stable cell line expressing FLAG-PKD2 was established in a similar manner with pCX4-FLAG-PKD2-IRES-Bsr by selecting cells with 20 μg/ml blasticidin S. A MDCK stable cell line expressing PAUF-MycHis was established in a similar manner with pMXs-PAUF-MycHis-IRES-Puro and 2 µg/ml puromycin. HeLa Tet-On cells were established with pRetroX-Tet-On Advanced and 500 µg/ml G418. HeLa Tet-On HA-mEGFP cells were established from the HeLa Tet-On cells with pRetroX-Tight-HA-mEGFP-Pur, 500 µg/ml G418, and 1 μg/ml puromycin. For establishment of a HeLa stable cell line expressing PAUF-MycHis, HeLa cells were transfected with pCI-PAUF-MycHis-IRES-Bsr and subjected to single-cell cloning with 5 μg/ml blasticidin S. All the other stable cell lines were obtained without single-cell cloning.

### Immunofluorescence microscopy

HeLa and MDCK cells were fixed with 4% paraformaldehyde (PFA) in phosphate-buffered saline (PBS) at room temperature for 20 min, permeabilized with 0.2% Triton X-100 in PBS for 30 min, and then blocked with 2% bovine serum albumin (BSA) in PBS for 30 min. The cells were labeled with the indicated primary antibodies and secondary antibodies conjugated to Alexa Fluor 350, 488, 594, or 647 in the blocking buffer. For specific staining of extracellularly located mKate2, HA-HeLa and HA-MDCK cells were fixed with 4% PFA and then blocked with 2% BSA without permeabilization, followed by labeling with an anti-mKate2 antibody and secondary antibodies conjugated to Alexa Fluor 488. The samples were analyzed with an Olympus Fluoview FV1200 confocal microscope with a UPLSAPO 60x/1.35 Oil objective and FV10-ASW software, an Olympus Fluoview FV3000 confocal microscope with a UPLXAPO 60x/1.42 Oil objective and FV31S-SW software, or an Evident Fluoview FV4000 confocal microscope with UPLXAPO 60x/1.42 and UPLXAPO 100x/1.45 Oil objectives and cellSens FV software. Image processing was performed with ImageJ software. Vesicular structures containing mKate2-FM4-HA/PAUF, EQ-SM, or PAUF-MycHis were detected using the plugin ComDet (https://github.com/UU-cellbiology/ComDet.git).

### Three-dimensional super-resolution microscopy

Super-resolution Z-stack images were acquired using the FV-OSR imaging system on an Evident Fluoview FV4000 confocal microscope with a UPLXAPO 100x/1.45 Oil objective and cellSens FV software. This system combines an optimized reduced confocal aperture with spatial frequency enhancement to achieve improved lateral resolution. Maximum intensity and 3D projections of the Z-stack images were generated using ImageJ software.

### Induction of HA-mEGFP expression by the Tet-On system

HeLa Tet-On HA-mEGFP cells were transfected with control or PKD2 siRNA. At 48 h after siRNA transfection, the medium was replaced with DMEM containing 10% FBS and 1 μg/ml doxycycline to induce expression of HA-mEGFP. After 24 h, the cells were fixed with 4% PFA and processed for immunofluorescence microscopy as described above.

### HA transport assay in HeLa cells

HA-HeLa cells were transfected with control siRNA or siRNA oligos targeting components of ER–Golgi MCSs, PKD2/3, or CAV1/2. At 48 h after siRNA transfection, the medium was replaced with DMEM containing 10% FBS; 24 h later, the medium was replaced with DMEM containing 10% FBS, 1 µM D/D solubilizer, and 20 µg/ml cycloheximide, and then the cells were incubated at 37°C for the indicated times. The cells were then fixed with 4% PFA and analyzed by fluorescence microscopy as described above.

### HA carrier and CARTS formation assays

For an HA carrier formation assay, HA-HeLa cells were transfected with control siRNA or siRNA oligos targeting components of ER–Golgi MCSs, PKD2/3, or CAV1/2. At 48 h after siRNA transfection, the medium was replaced with DMEM containing 10% FBS; 24 h later, the medium was replaced with DMEM containing 10% FBS, 20 mM Hepes-KOH, pH 7.4, 1 µM D/D solubilizer, and 20 µg/ml cycloheximide, and then the cells were incubated in a water bath at 20°C for 45min. The cells were then incubated in a water bath at 37°C for 5 or 15 min, followed by fixation with 4% PFA. The samples were analyzed by fluorescence microscopy as described above. A CARTS formation assay (Wakana & Tagaya, 2023) was performed in a similar manner with PAUF-HeLa cells. For experiments with OSW-1 or CRT0066101, HA-HeLa and PAUF-HeLa cells were incubated at 20°C for 45 min, pretreated with DMSO (control), 20 nM OSW-1, or 5 μM CRT0066101 for 15 min, and then shifted to 37°C for 5 or 15 min, followed by fixation with 4% PFA. Quantification of HA carriers and CARTS was performed with ImageJ software. The fluorescence signal of mKate2-FM4-HA/PAUF–containing puncta was distinguished from the background by setting a threshold and was analyzed at a set size within 0.05–2.00 µm^2^. For live-cell imaging, HA-HeLa cells were incubated with Opti-MEM containing 1 µM D/D solubilizer and 20 µg/ml cycloheximide in a water bath at 20°C for 45 min. The cells were then incubated at 37°C to monitor HA carrier biogenesis. Images were acquired continuously with a time interval between frames of 30 s for ∼90 min by use of an Olympus Fluoview FV1200 confocal microscope with a UPLSAPO 60×/1.35 Oil objective and FV10-ASW software. The images were processed with ImageJ software.

### HA transport assay in polarized MDCK cells

HA-MDCK cells were seeded onto Transwell inserts for polarization. After 96 h, the medium was replaced with DMEM containing 10% FBS, 1 μM D/D solubilizer, and 20 μg/mL cycloheximide, and then the cells were incubated at 37°C for the indicated times. For experiments with CRT0066101, the medium was replaced with DMEM containing 10% FBS, 20 mM HEPES-KOH, pH 7.4, 1 μM D/D solubilizer, and 20 μg/mL cycloheximide, and then the cells were incubated in a water bath at 20°C for 45 min, followed by pretreatment with DMSO (control) or 20 μM CRT0066101 for 15 min. The cells were then incubated in a water bath at 37°C for 5 or 30 min. The cells were then fixed with 4% PFA on the Transwell filters and processed for immunofluorescence microscopy as described above. The filters were cut out from the Transwell inserts and mounted on glass slides. The formation of tight junctions labeled with an anti-ZO-1 antibody confirmed the successful and uniform polarization of MDCK cells.

### PAUF secretion assay in polarized MDCK cells

MDCK cells stably expressing PAUF-MycHis were seeded onto Transwell inserts for polarization. After 96 h, the medium was replaced with Opti-MEM containing DMSO or 20 μM CRT0066101, and then the cells were incubated at 37°C for 15 min. After the replacement of the medium with fresh Opti-MEM containing DMSO or 20 μM CRT0066101, the cells were incubated at 37°C for 3 h. The medium was collected from both the upper (apical) and lower (basolateral) compartments, and the cells were lysed with 0.5% SDS and 0.025 U/µl Benzonase nuclease in PBS. Secreted proteins in the medium were precipitated with trichloroacetic acid, and the precipitates and cell lysates were analyzed by Western blotting with an anti-Penta-His antibody.

### CAV1 transport assay

HeLa cells were transfected with a plasmid for Str-Ii_CAV1-SBP-EGFP; 24 h later, the medium was replaced with DMEM containing 10% FBS, 1 µM D/D solubilizer, 20 µg/ml cycloheximide, and 40 µM biotin or 10 µM BME, and then the cells were incubated at 37°C for the indicated times. The cells were then fixed with 4% PFA and analyzed by fluorescence microscopy as described above. In cotransport assays of CAV1 and HA, or CAV1 and PAUF, HA-HeLa and PAUF-HeLa cells were transfected with a plasmid for Str-Ii_CAV1-SBP-EGFP; 24 h later, the medium was replaced with DMEM containing 10% FBS, 1 µM D/D solubilizer, and 20 µg/ml cycloheximide, and then the cells were incubated at 37°C; 5 min later, BME was added to a final concentration of 10 µM, and then the cells were incubated at 37°C for 25 min, followed by fixation with 4% PFA. For live-cell imaging, PAUF-HeLa cells were transfected with a plasmid for Str-Ii_CAV1-SBP-EGFP; 24 h later the medium was replaced with Opti-MEM containing 1 µM D/D solubilizer, 20 µg/ml cycloheximide, and 10 µM BME, and the cells were incubated in a water bath at 20°C for 35 min. The cells were then incubated at 37°C to monitor TGN export of CAV1-SBP-EGFP and mKate2-FM4-PAUF. Images were acquired continuously with a time interval between frames of 3 s for ∼3.5 min by use of an Evident IXplore IX85 Spin confocal spinning disk microscope with a UPLXAPO 100x/1.45 Oil objective and cellSens Dimension software. The images were processed with ImageJ software.

### CD-M6PR transport assay

HeLa cells were transfected with control siRNA or siRNA oligos targeting CAV1/2. At 48 h after siRNA transfection, the cells were transfected with a plasmid for Str-KDEL_SBP-EGFP-CD-M6PR; 24 h later, the medium was replaced with DMEM containing 10% FBS, 40 µM biotin, and 20 µg/ml cycloheximide, and then the cells were incubated at 37°C for the indicated times. The cells were then fixed with 4% PFA and analyzed by fluorescence microscopy as described above.

### Statistical analysis

Data analysis was performed using GraphPad PRISM software. The graph data shown are for a single representative experiment out of at least three performed, unless otherwise stated.

### Quantification of HA/PAUF incorporation into TGN-derived tubules

HA/PAUF-HeLa cells containing mKate2-FM4-HA/PAUF–positive TGN-derived tubules were counted manually using Fiji. For quantification of HA/PAUF incorporation into PKD2 KD-induced membrane tubules, the tubules were defined as GST-PKD2 KD-positive tubular structures emerging from the perinuclear region or otherwise clearly distinguishable. The total number of PKD2 KD-positive membrane tubules per cell was counted, and the proportion of tubules containing mKate2-FM4-HA or mKate2-FM4-PAUF was determined. Quantifications were performed independently twice in a blinded manner, and the mean values were used for statistical analysis. Blinding was achieved using a custom Fiji macro that randomly renamed all images and generated a separate CSV mapping file linking each randomized image to its original file; this map was used only after quantification to decode the results.

### Quantification of fluorescence intensity in the Golgi complex and the PM

Relative HA fluorescence intensity quantification between the PM and the Golgi complex was performed using a custom-written MATLAB pipeline available at https://github.com/JessicaAngulo/Hemagglutinin_secretion.git. For HeLa cells, whole-cell segmentation was carried out manually using the *drawpolygon* function in MATLAB. For MDCK cells, segmentation was performed automatically using the fluorescence microscopy channel of the tight junction marker ZO-1. Images of MDCK cells were pre-processed by applying a Wiener filter (*wiener2* function, using a 10×10 window) to reduce noise, followed by top-hat filtering with a disk-shaped structuring element (radius 15) using *imtophat* function, to enhance bright features. Local minima of less than 3000 grey values (16 bits images) were suppressed using the *imhmin* function, and final segmentation was performed via the *watershed* algorithm. This step also allowed exclusion of out-of-focus cells. To segment the Golgi area, a fixed intensity threshold was applied to the respective Golgi marker channel (120/256 grey levels on the TGN46 channel for HeLa Tet-On HA-mEGFP cells, and 70/256 grey levels on the GM130 channel for HA-MDCK cells or HA-HeLa cells expressing each CAV1-mEGFP construct), using images acquired at the z-plane that corresponds to the equatorial plane of the Golgi-positive signal. Thresholded images were contrast-adjusted with *imadjust* and Gaussian-filtered with a standard deviation of 2 with *imgaussfilt*. Binary masks of the Golgi regions were generated and refined using *bwboundaries* with the ‘noholes’ option to identify structure boundaries. Quantification of HA fluorescence intensity density was performed by integrating the signal over each segmented region of interest (ROI) –either the entire PM mask or the Golgi mask– and normalizing by the corresponding ROI area (whole-cell area or Golgi-positive area, respectively). For the microscopy images acquired with a single confocal plane, as exemplified in Fig. 6A, the fluorescence intensity in the PM was defined and quantified as the fluorescence intensity measured in the single plane within the entire manually segmented cell mask minus the fluorescence intensity in the segmented Golgi area. The PM/Golgi HA intensity ratio was then calculated by dividing the two intensity densities and subsequently normalized to the corresponding control condition.

### Image analysis for characterization of SBP-EGFP-CD-M6PR–positive puncta

The analysis was performed using a custom-written MATLAB pipeline available at https://github.com/JessicaAngulo/Carrier_characterization.git. Whole-cell segmentation was carried out manually using the *drawpolygon* function in MATLAB. A global threshold for carrier detection was determined by applying a Gaussian filter followed by morphological erosion to enhance vesicle separation. A pixel intensity threshold was set based on the image mean plus one standard deviation. The same threshold was applied across all images within a condition to ensure consistency. After thresholding, vesicular structures were identified using binary masks and bwboundaries. Only structures with areas between 5 and 500 pixels were retained. For each carrier, the area was calculated.

### Analysis of CAV1 enrichment in PKD2 KD-positive bulges

Images were analyzed using a custom Fiji macro to segment PKD2 KD-positive structures and quantify CAV1 fluorescence. Individual cells were manually outlined using the freehand selection tool. The PKD2 KD channel was duplicated and restricted to the selected cell area, after which a Gaussian filter (sigma = 2.0) was applied. PKD2 KD-positive puncta were then segmented using the automatic Yen threshold, which was visually inspected and minimally adjusted when necessary to ensure appropriate visual correspondence with the punctate signal. The resulting binary mask was subjected to watershed segmentation, and puncta were identified using particle analysis with an area range of 0.15–5.0 µm² and a minimum circularity of 0.50. For each detected PKD2 KD-positive particle, CAV1 fluorescence intensity was measured and normalized to the mean CAV1 fluorescence intensity of the corresponding whole-cell area. Histograms of the normalized relative intensities were generated, and puncta with a >5-fold increase in CAV1 relative intensity were classified as CAV1-enriched.

### Immunoprecipitation

HEK 293T cells stably expressing FLAG-PKD2 WT were transfected with a plasmid for EGFP (control) or CAV1 WT-mEGFP; 24 h later, HEK 293T cells were washed with PBS and lysed in buffer A (50 mM Tris-HCl, pH 7.4, 150 mM NaCl, 0.1% SDS, 0.5% sodium deoxycholate, 1% Triton X-100, 1 µg/ml leupeptin, 2 µM pepstatin A, 2 µg/ml aprotinin, and 1 mM phenylmethylsulfonyl fluoride), and the lysates were centrifuged at 17,700× g for 10 min. The supernatants were further centrifuged at 100,000× g for 30 min, and the resulting supernatants were immunoprecipitated with GFP-Trap Agarose. After incubation at 4°C for 2 h, the beads were washed four times with buffer B (buffer A without protease inhibitors), and the precipitated proteins were then eluted with SDS sample buffer and analyzed by Western blotting using anti-FLAG and anti-GFP antibodies, followed by horseradish peroxidase-conjugated secondary antibodies.

### GST pull-down

HEK 293T cells were transfected with a plasmid for GST (control) or GST-PKD2 (WT or KD) in combination with a plasmid for CAV1 (WT or P132L)-mEGFP; 24 h later, the cells were washed with PBS and lysed in buffer A. The lysates were centrifuged at 17,700× g for 10 min and the resulting supernatants were mixed with Glutathione Sepharose 4B. After incubation at 4°C for 2 h, the beads were washed four times with buffer B, and the bound proteins were then eluted with SDS sample buffer and analyzed by Western blotting using anti-GFP and anti-GST antibodies, followed by horseradish peroxidase-conjugated secondary antibodies.

### BiFC analysis

HeLa Tet-On cells were cotransfected with pCX4-Vn-PKD2-IRES-Bsr and pRetroX-Tight-CAV1-Vc-Pur in the presence of 1 μg/ml doxycycline. After 24 h, the cells were fixed with 4% PFA and processed for immunofluorescence microscopy as described above. Co-staining of the cells with anti-Vn and anti-Vc antibodies showed that nearly 100% of Vn-PKD2–expressing cells co-expressed CAV1-Vc. To quantify BiFC signal at the TGN, confocal images of the cells co-stained with anti-Vn and anti-TGN46 antibodies were analyzed using Fiji/ImageJ software. A TGN mask was generated from the TGN46 channel by applying a 1-pixel radius Gaussian smoothing, automatic thresholding, and a single dilation step to create a binary mask encompassing the TGN region. The BiFC fluorescence channel was multiplied by the TGN46-derived binary mask to specifically restrict BiFC signal within the TGN region. For each cell displaying detectable BiFC signal, the total BiFC fluorescence intensity within the TGN46-defined mask was measured, thereby minimizing variability arising from differences in TGN size or morphology. To reduce the impact of cell-to-cell variability in expression levels, BiFC values were normalized on a per-experiment basis: for each experimental day, values from all quantified cells were normalized to the median BiFC signal of the PKD2-WT+CAV1-WT condition.

## Data availability

The raw image data generated in this study have been deposited in the BioImage Archive under accession number S-BIAD4143 and will be made publicly available upon publication.

## Acknowledgments

We thank Yoshihiro Mimaki, Sang Seok Koh, Hans-Peter Hauri, Salvatore Chiantia, Juan Bonifacino, Franck Perez, Ambra Pozzi, Ari Helenius, Francis A. Barr, Christopher G. Burd, and Vivek Malhotra for providing materials. We appreciate the technical assistance of Wataru Goto, Riko Kazama, Itsuki Minowa, Aoi Sakaguchi, Ayui Mizutani, Tatsuki Saito, Yuho Masamoto, Sora Ito, and Hiyori Makabe. This work was supported in part by Grants-in-Aid for Scientific Research from the Ministry of Education, Culture, Sports, Science, and Technology of Japan (grant numbers 20K06562 and 25K09568 to Y. Wakana), AMED Multidisciplinary Frontier Brain and Neuroscience Discoveries (Brain/MINDS 2.0) (grant number JP24wm0625506 to Y. Wakana), the Takeda Science Foundation (to Y. Wakana), Daiichi Sankyo Foundation of Life Science (to Y. Wakana), and the Ono Medical Research Foundation (to Y. Wakana). F. Campelo and J. Angulo-Capel acknowledge support by the Government of Spain (CEX2024-001490-S, MICIU/AEI/10.13039/501100011033), State Research Agency (AEI) PID2022−138282NB-I00 project funded by the MCIN/AEI/10.13039/501100011033/FEDER, UE, “Unidad de Excelencia María de Maeztu” CEX2024-001431-M, funded by MICIU/AEI/10.13039/501100011033 to MELIS-UPF, Fundació CELLEX (Barcelona), Fundació Privada Mir-Puig, the Generalitat de Catalunya (CERCA, AGAUR). F. Campelo is also supported by “Unidad de Excelencia María de Maeztu” CEX2024-001431-M, funded by MICIU/AEI/10.13039/501100011033. This project has received funding from the European Union’s Horizon 2020 research and innovation programme under Marie Skłodowska-Curie grant agreement number 847517 (to J. Angulo-Capel). E. Almacellas acknowledges the support from the Beatriu de Pinós postdoctoral fellowship program of the Department of Research and Universities of the Generalitat de Catalunya (2023 BP 00210) and from the European Union’s Horizon Europe research and innovation programme under the Marie Sklodowska-Curie grant agreement number 101205472.

## Expanded view figure legends

**Figure EV1.**
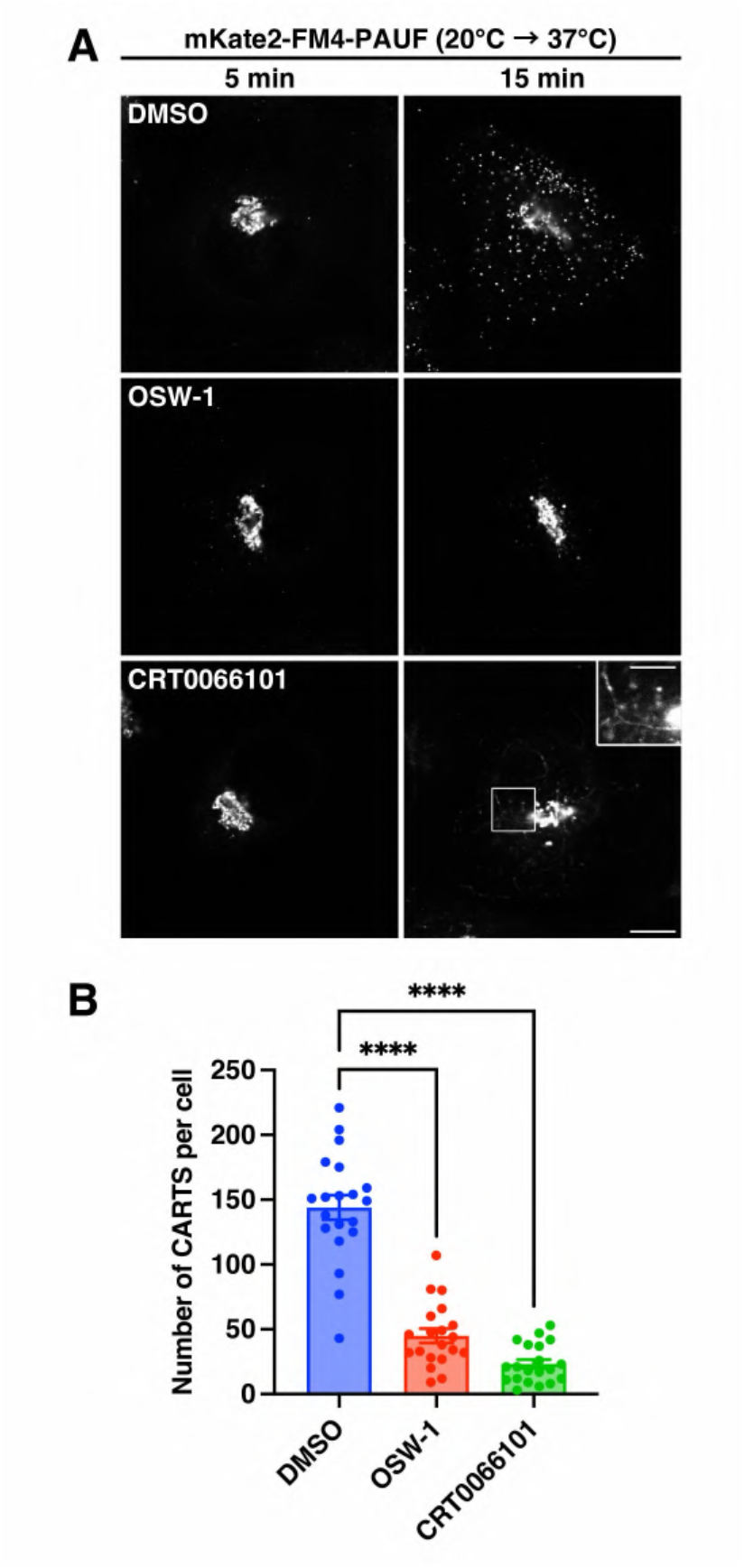
Inhibitory effects of OSW-1 and CRT0066101 on CARTS biogenesis. **(A, B)** CARTS biogenesis in HeLa cells stably expressing mKate2-FM4-PAUF (PAUF-HeLa cells) treated with DMSO, 20 nM OSW-1, or 5 μM CRT0066101. The cells were subjected to CARTS formation assay as described in Methods, fixed, and visualized by mKate2 fluorescence. The inset shows a magnified image of the boxed area with brightness and contrast enhancement. Scale bars, 10 µm (large panels) and 5 µm (inset). **(B)** Quantification of the CARTS biogenesis. The number of CARTS (mKate2-FM4-PAUF–positive dots) at 15 min after the temperature shift to 37°C is shown. Data are means ± SEM (n = 20 cells per condition; ****, P < 0.0001; one-way ANOVA multiple comparison test). Source data are available online for this figure.

**Figure EV2.**
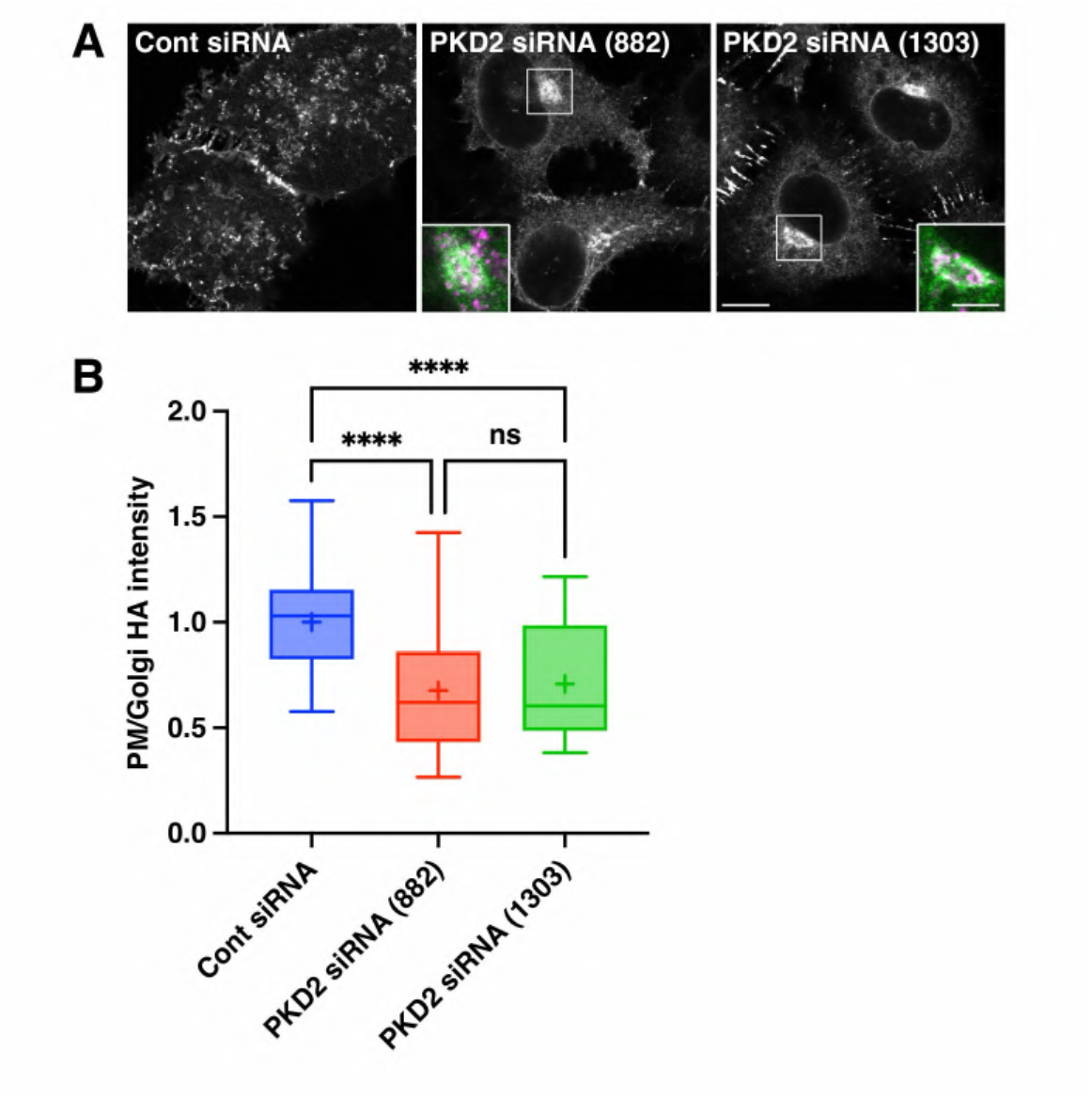
Effects of PKD2 knockdown on distribution of full-length HA-mEGFP. **(A, B)** Accumulation of HA-mEGFP at the Golgi complex under PKD2 knockdown conditions in HeLa Tet-On HA-mEGFP cells. The cells were fixed and visualized by mEGFP fluorescence and anti-TGN46 immunostaining. The insets show magnified and merged images of HA-mEGFP (green) and the TGN maker TGN46 (magenta) in the boxed areas. Scale bars, 10 µm (large panels) and 5 µm (insets). **(B)** Quantification of HA-mEGFP distribution between the PM and the Golgi complex. The ratio of fluorescence intensity of HA-mEGFP in the PM and the Golgi complex, normalized as the values in control cells, is shown. Boxes delimit the first and third quartiles, and the central line is the median, whereas the cross represents the mean value. The whiskers represent the minimum and maximum values (control [Cont] siRNA: n = 95 cells; PKD2 siRNA [882]: n = 90 cells; PKD2 siRNA [1303]: n = 78 cells; ****, P < 0.0001; one-way ANOVA multiple comparison test). Source data are available online for this figure.

**Figure EV3.**
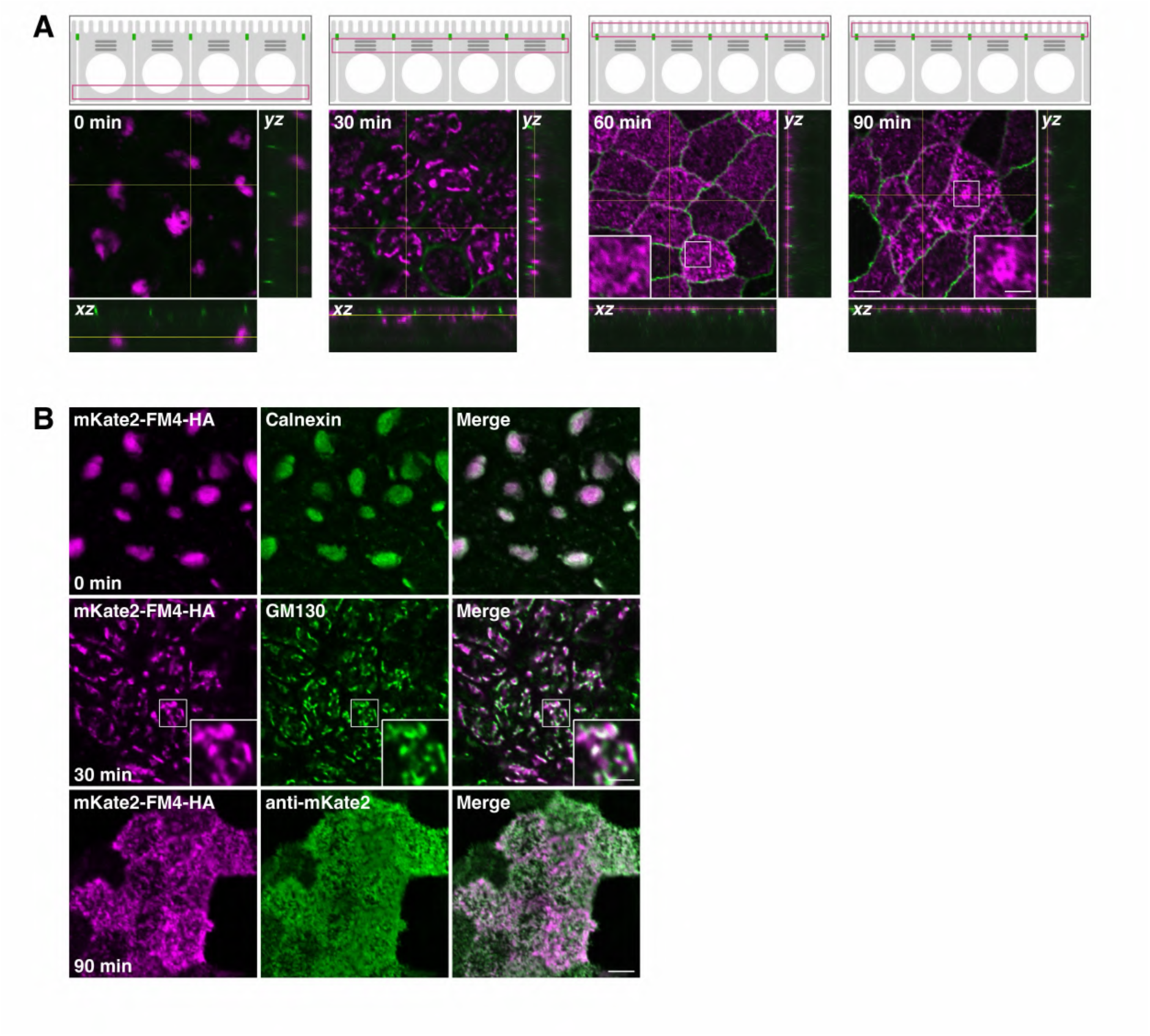
Synchronized transport of HA in polarized MDCK cells. **(A)** mKate2-FM4-HA transport from the ER to the apical PM via the Golgi complex in polarized HA-MDCK cells. HA-MDCK cells were incubated at 37°C with D/D solubilizer and cycloheximide, fixed at the indicated times, and visualized by mKate2 fluorescence and anti-ZO-1 immunostaining. Merged images of mKate2-FM4-HA (magenta) and the tight junction marker ZO-1 (green) are shown. Each schematic representation at the top indicates the location corresponding to the largest panel. **(B)** Colocalization of mKate2-FM4-HA with the ER maker calnexin (0 min) and the Golgi maker GM130 (30 min) in permeabilized HA-MDCK cells and visualization of extracellularly located mKate2 (90 min) by labelling of non-permeabilized HA-MDCK cells with an anti-mKate2 antibody. The insets show magnified images of the boxed area. Scale bars, 5 µm (large panels) and 2 µm (insets). Source data are available online for this figure.

**Figure EV4.**
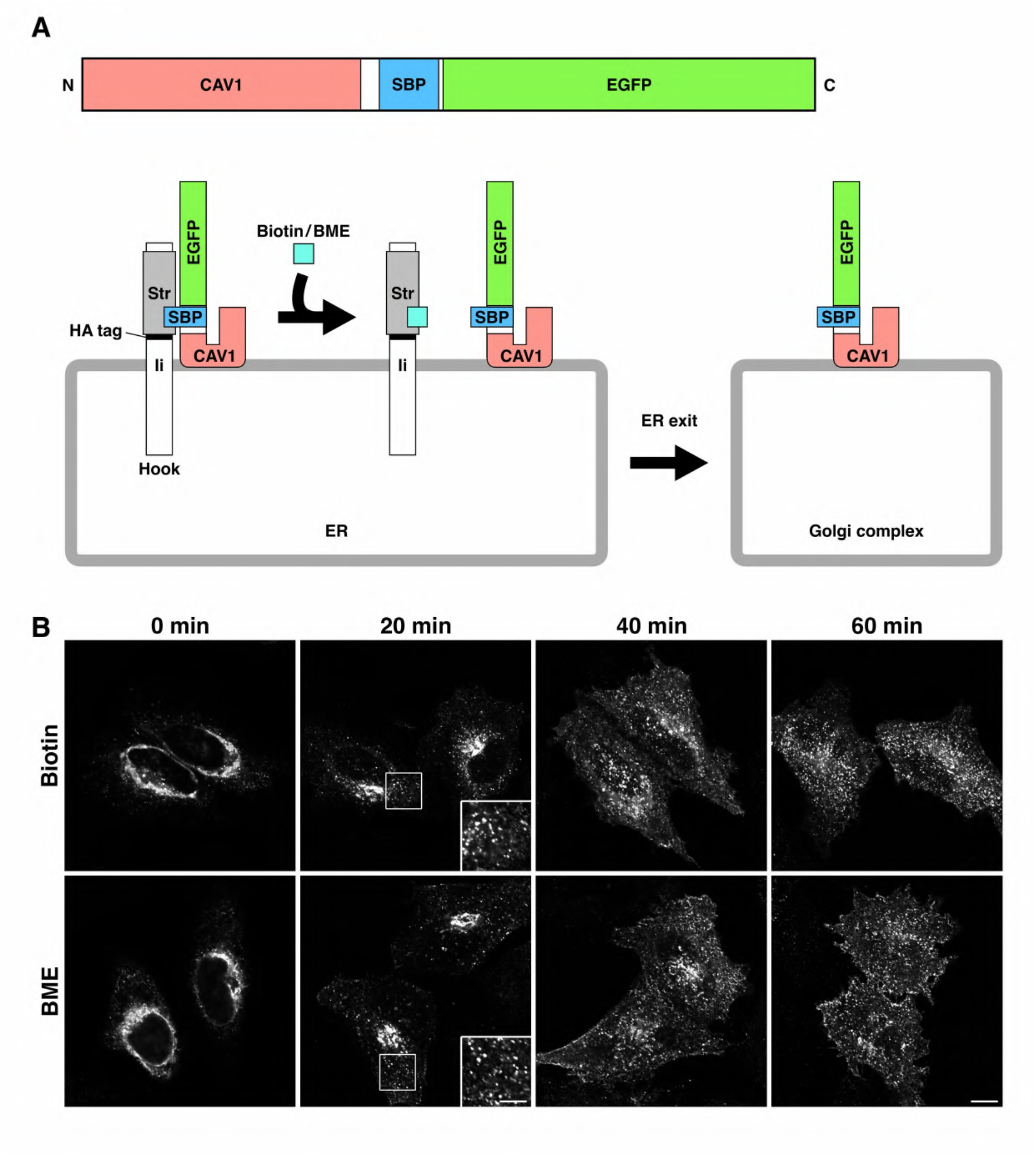
Synchronized transport of CAV1. **(A)** Schematic representations of the CAV1-SBP-EGFP construct and its synchronized transport from the ER to the Golgi complex by using the RUSH system. **(B)** CAV1-SBP-EGFP transport from the ER to the PM via the Golgi complex in HeLa cells. HeLa cells expressing Str-Ii_CAV1-SBP-EGFP were incubated at 37°C with biotin (upper panels) or BME (lower panels), fixed at the indicated times, and visualized by EGFP fluorescence. The insets show magnified images of the boxed areas with brightness and contrast enhancement. Scale bars, 10 µm (large panels) and 5 µm (insets). Source data are available online for this figure.

**Figure EV5.**
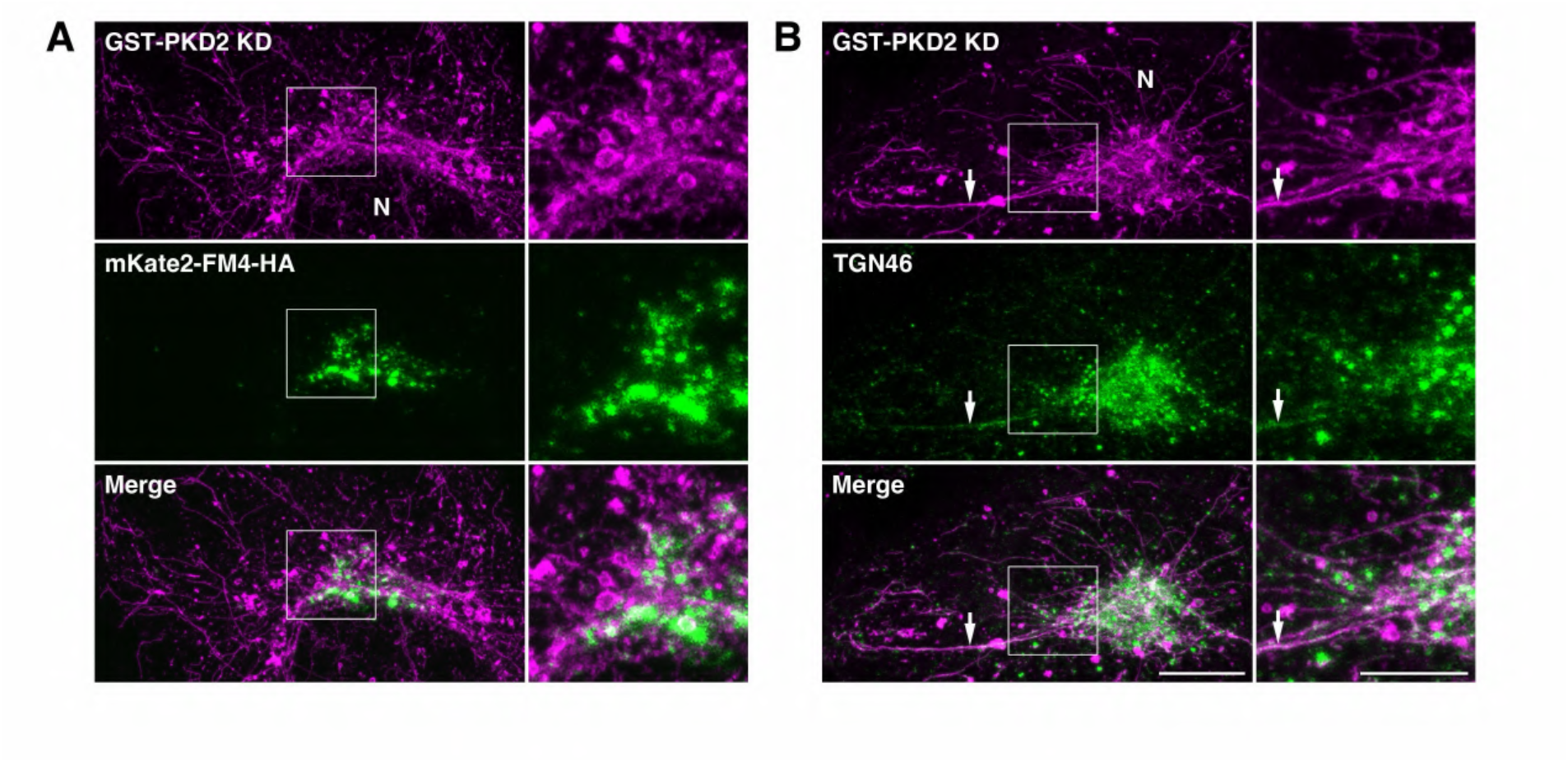
Distribution of mKate2-FM4-HA and TGN46 upon GST-PKD2 KD expression. **(A)** 3D super-resolution microscopic images of GST-PKD2 KD and mKate2-FM4-HA at 30 min after transport initiation. HA-HeLa cells expressing GST-PKD2 KD were incubated at 37°C for 30 min with D/D solubilizer and cycloheximide, fixed, and visualized by anti-GST immunostaining and mKate2 fluorescence. **(B)** 3D super-resolution microscopic images of GST-PKD2 KD and TGN46. HeLa cells expressing GST-PKD2 KD were fixed and subjected to anti-GST and anti-TGN46 immunostaining. Maximum intensity merges of Z-stack images are shown with magnified views of the boxed areas (right of each panel). Arrows indicate GST-PKD2 KD- and TGN46-containing long tubules extended from the TGN. N, nucleus. Scale bars, 10 µm (left panels) and 5 µm (right panels). Source data are available online for this figure.

**Movie EV1. HA carrier biogenesis at the TGN.** HA-HeLa cells were first incubated at 20°C with D/D solubilizer and cycloheximide for 45 min. Images were acquired continuously after the temperature shift to 37°C with a time interval between frames of 30 s for ∼90 min by use of an Olympus Fluoview FV1200 confocal microscope. Scale bar, 10 µm.

**Movie EV2. Co-export of CAV1-SBP-EGFP (green) and mKate2-FM4-PAUF (magenta) from the TGN.** PAUF-HeLa cells expressing Str-Ii_CAV1-SBP-EGFP were first incubated at 20°C with D/D solubilizer, cycloheximide, and 10 µM BME for 35 min. Images were acquired continuously after the temperature shift to 37°C with a time interval between frames of 3 s for ∼3.5 min by use of an Evident IXplore IX85 Spin confocal spinning disk microscope. CAV1-SBP-EGFP, green; mKate2-FM4-PAUF, magenta. Scale bar, 10 µm.

**Movie EV3. CAV1-positive tubule formation during CARTS biogenesis.** An enlarged portion of Movie EV2 is shown. CAV1-SBP-EGFP, green; mKate2-FM4-PAUF, magenta. Scale bar, 1 µm. See also Appendix Fig. S2.

**Movie EV4. Accumulation of CAV1 WT-mEGFP at bulging TGN domains connected to GST-PKD2 KD-positive tubules.** A 3D projection of Z-stack super-resolution microscopic images of HeLa cells expressing GST-PKD2 KD (magenta) and CAV1 WT-mEGFP (green) is shown. Scale bar, 10 µm.

