## Appendix for "A protein kinase D–caveolin axis drives secretory carrier biogenesis at the trans-Golgi network"

### Table of Contents

Appendix Figure S1 Page 2
Appendix Figure S2 Page 3
Appendix Figure S3 Page 4
Appendix Table S1 Page 5

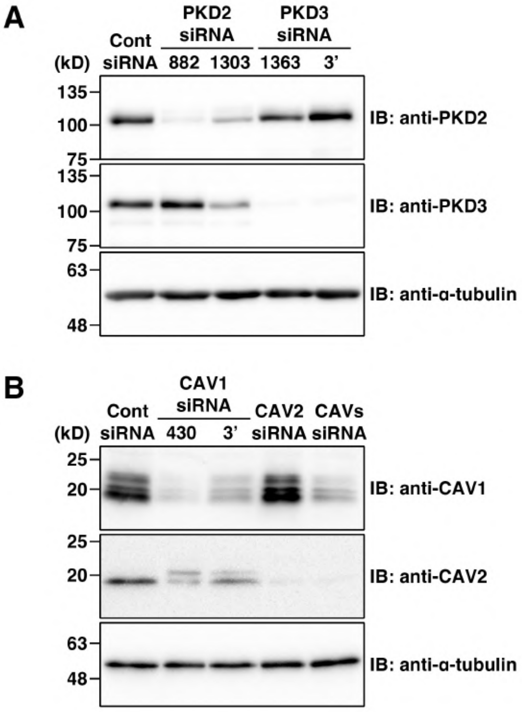


**Appendix Figure S1. siRNA-mediated depletion of PKD and CAV isoforms.**
(**A**) Knockdown efficiency of PKD2 and PKD3 in HeLa cells at 72 h after siRNA transfection. The cell lysates were immunoblotted (IB) with the indicated antibodies. (**B**) Knockdown efficiency of CAV1 and CAV2 in HeLa cells at 72 h after siRNA transfection. Multiple bands in the immunoblot with an anti-CAV1 antibody represent CAV1α, CAV1β, and their modified forms. As a loading control, α-tubulin was detected in (**A**, **B**).

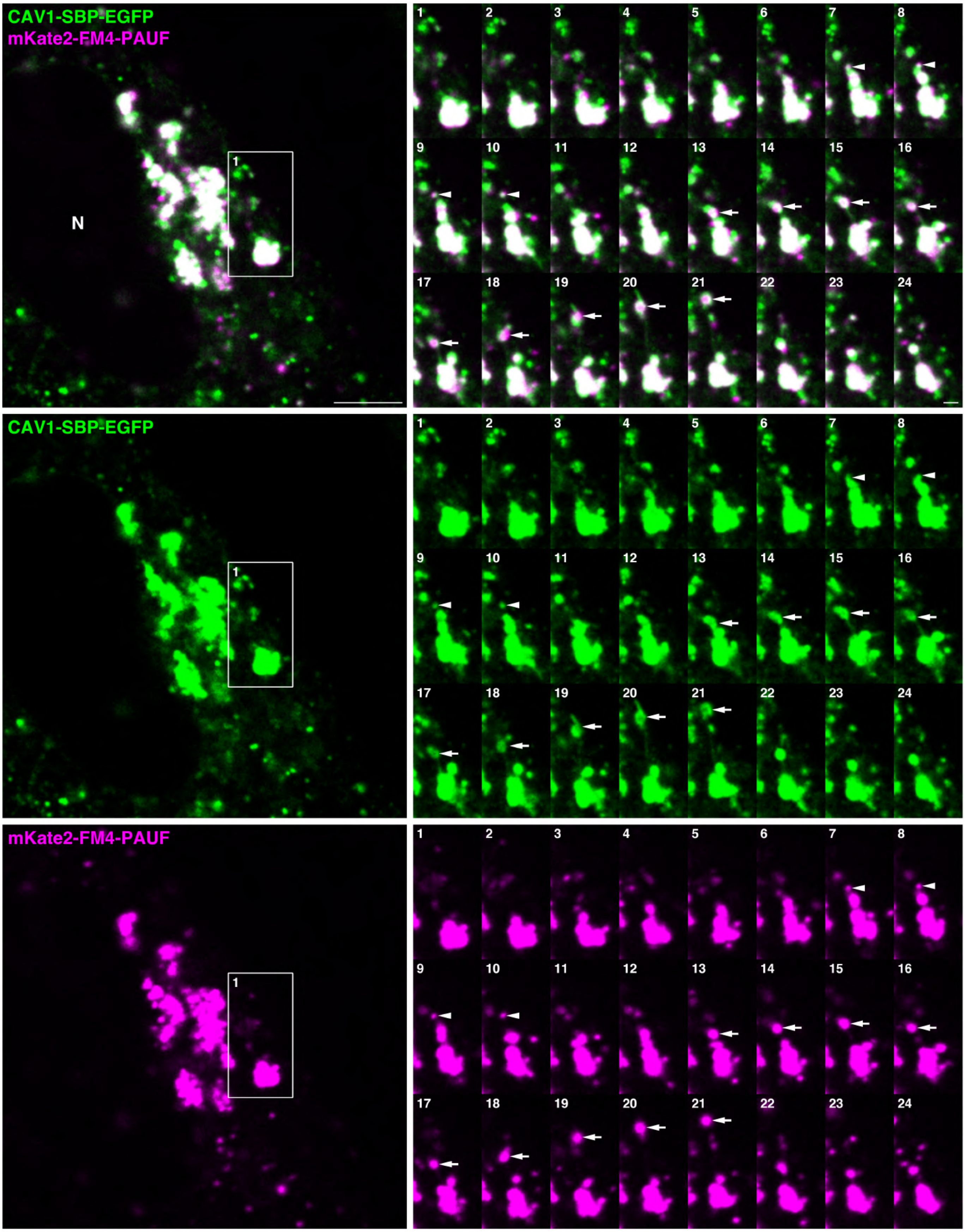


**Appendix Figure S2. CAV1-positive tubule formation during CARTS biogenesis.**
The left panels show the 26th frame of Movie EV2. The boxed region indicates the area enlarged in Movie EV3. The right panels show the complete time series of Movie EV3. Merged images are shown in the top row, with the corresponding individual channels shown below. Arrowheads indicate an mKate2-FM4-PAUF–positive CARTS carrying CAV1-SBP-EGFP, whereas arrows indicate a larger mKate2-FM4-PAUF–positive CARTS associated with surface-localized CAV1-SBP-EGFP and connected to CAV1-SBP-EGFP–positive tubules. Scale bars, 5 μm (left panels) and 1 μm (right panels).

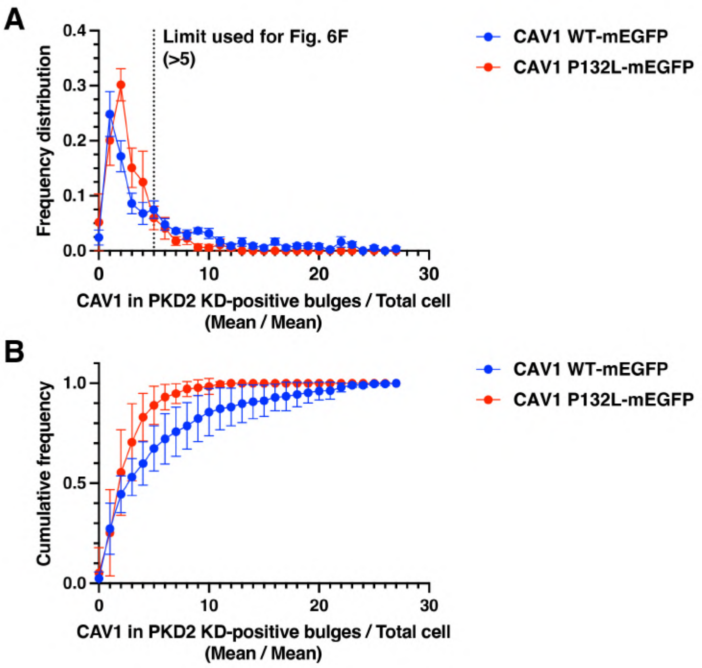


**Appendix Figure S3. Quantification of CAV1 enrichment at PKD2 KD-induced TGN bulging domains.** (**A**) Frequency distribution of the mean fluorescence intensity of CAV1 (WT or P132L)-mEGFP, normalized as described in Methods. The dotted line indicates the >5-fold enrichment threshold used for the quantification shown in Fig. 6F. (**B**) Cumulative frequency distribution corresponding to the data shown in (**A**).

**Appendix Table S1. Targeting sequences of siRNA oligos used in this study**

| Identifier | Sequence | Source |
|---|---|---|
| Control (GL2 luciferase) | 5'-AACGTACGCGGAATACTTCGA-3' | Wakana *et al*, 2015, 2021 |
| VAP-A | 5'-AACTAATGGAAGAGTGTAAAA-3' | Wakana *et al*, 2015, 2021 |
| VAP-B | 5'-AAGAAGGTTATGGAAGAATGT-3' | Wakana *et al*, 2015, 2021 |
| OSBP | 5'-AAGGAGTTAACCCATATTTAT-3' | This study |
| Sac1 | 5'-AACTGATATTCAGTTACAAGA-3' | Wakana *et al*, 2021 |
| SCAP | 5'-AACCTCCTGGCAGTAGATGTA-3' | Wakana *et al*, 2021 |
| CERT | 5'-AACATTCTATGCAAGATTACA-3' | This study |
| PKD2 (882) | 5'-ATGCAAAGACTGCAAGTTTAA-3' | This study |
| PKD2 (1303) | 5'-AACAGATACTATAAGGAAATT-3' | This study |
| PKD3 (1363) | 5'-AAGTATTATAAGGAAATTCCA-3' | This study |
| PKD3 (3') | 5'-AAGATATGAAGAAATATGATA-3' | This study |
| CAV1 (430) | 5'- CAGAAAGAAATATAAATGACA-3' | This study |
| CAV1 (3') | 5'-AAGGTAATTTGAGAGAAATAT-3' | This study |
| CAV2 | 5'-AACCACATTTAGAAATGTTTA-3' | This study |
